# Asymmetric Adaptation of Catch-up Saccades with Artificial Central Scotomas

**DOI:** 10.64898/2026.09.28.755127

**Authors:** Sydney Doré, Jenna Black, Gunnar Blohm, Aarlenne Z Khan

**Affiliations:** Queen’s University, Kingston, ON, Canada; School of Optometry, Université de Montréal, Montréal, Canada

**Keywords:** Saccade, Pursuit, Catch-up Saccade, Scotoma, Age-related Macular Degeneration, Central Vision Loss, Adaptation

## Abstract

**Purpose:** In fully-sighted individuals, pursuit and saccadic eye movements work together to foveate moving targets; pursuit attempts to match target velocity, while catch-up saccades re-align the fovea to the target if it deviates during pursuit. Although saccade-pursuit interactions are well understood in healthy individuals, less is known about how these mechanisms adapt in the presence of central vision loss.

**Methods:** We simulated central vision loss in healthy young adults to examine plasticity in saccade-pursuit mechanisms, investigating whether and how catch-up saccades adapted over five days of visual tracking. 25 fully-sighted participants performed a task where they pursued a moving target that stepped position randomly (designed to elicit catch-up saccades) while viewing with a gaze-contingent central scotoma. The scotoma was shaped like a bitten cookie to encourage use of a specific preferred retinal locus (PRL) corresponding to the peripheral location with the highest acuity.

**Results:** After five days of repetition, most participants’ catch-up saccade endpoints shifted towards the PRL rather than the fovea. On average, endpoints shifted 0.5° towards the PRL between days 1 and 5 (0.3°-1.5°), with significant shifts in 13/25 participants. Adaptation was asymmetric; shifts were largest when the target moved towards the PRL (into the scotoma; towards cartesian quadrant 1) and smallest away from the PRL (out of the scotoma; towards cartesian quadrant 3).

**Conclusions:** In summary, we observed adaptation of saccade-pursuit interactive mechanisms through re-referencing of catch-up saccades from fovea to PRL. These results provide a promising avenue for rehabilitation in central vision loss aimed at improving eye movements and encouraging PRL use. Ultimately, such an approach may enhance residual visual function and quality of life.

**Significance Statement:** Individuals with central vision loss can shift their oculomotor reference from the fovea to a preferred retinal locus (PRL) during saccades to stationary targets. Whether this extends to moving targets remains unclear. Our findings demonstrate catch-up saccade adaptation to a PRL after 5 hours of training. Our results support rehabilitation strategies promoting efficient PRL use, allowing individuals with central vision loss to optimize their residual vision.

## 1. Introduction

Fully sighted humans visually track moving objects by combining smooth pursuit and saccadic eye movements. Smooth pursuit is used to stabilize the image of a moving target on the fovea and is primarily driven by target motion, while saccades are rapid, ballistic movements driven by a position error between a visual target and the eye (Rashbass, 1961). The mechanisms of saccade-pursuit interactions are generally well understood, including how these two systems coordinate to maintain stable gaze on a moving target (Orban de Xivry et al., 2006; Orban de Xivry & Lefèvre, 2007). When smooth pursuit fails to match the speed of an object, a catch-up saccade is triggered to correct the tracking error. Importantly, these catch-up saccades use the fovea as the oculomotor reference point, meaning that they are planned such that the target lands on the fovea at saccade completion. Thus, the saccade and pursuit eye movement systems coordinate to ensure that the image of the moving object is maintained on the fovea, the region of highest visual acuity of the macula. However, in individuals with central vision loss, the fovea is non-functional, raising the question of how the planning of catch-up saccades adapt when the primary oculomotor reference is no longer available.

One of the leading causes of central visual loss is age-related macular degeneration (AMD). In macular degeneration, damage to foveal photoreceptors results in a central scotoma that impairs high-acuity vision in one or both eyes. Without functional foveal vision, patients must rely on peripheral vision, which has considerably lower spatial resolution and is poorly suited to tasks that normally depend on foveal processing. Central vision loss associated with AMD impairs the ability to perform daily tasks such as reading, recognizing faces and driving, significantly reducing patients’ quality of life. Despite central vision loss, individuals with AMD can still pursue moving objects using peripheral vision (González et al., 2018; Shanidze et al., 2016), though pursuit gain is reduced and catch-up saccades occur more frequently (González et al., 2018; Shanidze et al., 2016; Shanidze & Velisar, 2020; Verghese et al., 2021). A likely explanation for this degraded performance is that the oculomotor system continues to use the damaged fovea as its reference point, even in the presence of a scotoma, generating unnecessary and inaccurate catch-up saccades directed toward a region that can no longer provide reliable visual feedback and ultimately impairing tracking performance.

Some patients with central vision loss develop a preferred retinal locus (PRL), a peripheral retinal region used as an functional substitute to the fovea, typically located in the areas of highest remaining acuity (Crossland et al., 2005; Greenstein et al., 2008; White & Bedell, 1990). PRLs can develop spontaneously following bilateral foveal loss, typically emerging after six months and are used for tasks such as reading (Crossland et al., 2005; Verghese et al., 2021). Individuals can also develop multiple PRLs, switching between them across different tasks according to their specific visual demands (Shanidze et al., 2016). Spontaneously developed PRLs are not always optimally located for all tasks, however. Some studies suggest that individuals can be trained to adopt a more favorable retinal location, referred to as a trained retinal locus (TRL). For example, patients whose PRLs were poorly positioned for reading were successfully trained over five hours to use a TRL in a more favorable location, resulting in significant improvements in reading speed (Nilsson et al., 2003). While much of this work has focused on fixation tasks and eye movements from stationary targets, less is known about how the PRL is incorporated into the oculomotor system as a reference point for gaze.

A small number of studies have examined whether saccades can be planned and executed to align the target with the PRL rather than the fovea, with this shift occurring for some patients with macular degeneration either spontaneously or through explicit eccentric viewing training (Cheung & Legge, 2005; Heinen & Skavenski, 1992; White & Bedell, 1990; Whittaker et al., 1988). This has been shown primarily for stationary scenes or images. Oculomotor referencing has also been demonstrated in fully sighted individuals with simulated scotomas (Kwon et al., 2013; Léné et al., 2020). Under these simulated scotoma conditions, the oculomotor system has been shown to rapidly adopt a PRL and re-reference saccades from fixation in as little of 15 hours of training (Léné et al., 2020). It has also been shown that trained re-referencing to the PRL can be retained over weeks after training (Kwon et al., 2013). Taken together, these studies demonstrate that training can result in oculomotor re-referencing to the PRL for static targets in both real and simulated scotomas conditions. Whether this extends to catch-up saccades during pursuit of moving targets remains unknown, as does the question of whether behavioural training can establish or reinforce PRL use to improve visual tracking. Here, we examined plasticity in saccade-pursuit mechanisms, focusing on the catch-up saccade component of visual tracking by testing whether and how young, fully sighted participants adapted their visual tracking behaviour in the presence of a simulated central scotoma over the course of five days of training. We tested younger participants, who show greater plasticity than older adults (Léné et al., 2020) and are therefore the most likely group to exhibit a training-induced shift in the oculomotor reference for catch-up saccades from the fovea to the PRL.

## 2. Methods

### 2.1 Participants

Twenty-five fully sighted participants (9 males; mean age = 23 years, range = 19–35 years) with normal or corrected-to-normal vision took part in the experiment. The study’s procedures were approved by the Queen’s University General Research Ethics Board in accordance with the Declaration of Helsinki. Participants provided informed consent and received compensation for their participation.

### 2.2 Experiment Setup and Stimuli

Participants sat in a semi-dark room with their head stabilized on a chin/head rest viewing stimuli displayed on a ViewPixx screen (VPixx Technologies, 120 Hz refresh rate, resolution 1,920 × 1,200, strobed backlight) at 62 cm. Eye position of the right eye was recorded using an Eyelink 1000 video-based recording system (SR Research, Mississauga, ON, Canada) at 1,000 Hz with a tower-mount setup. At the beginning of each experimental block, a 2-step calibration process was undertaken to ensure that the gaze-contingent scotoma was aligned with gaze with high accuracy. Each participant first completed a standard 9-point Eyelink calibration, followed by a second calibration, in which they fixated on 30 dots, presented randomly at different locations on the screen. A 6-parameter polynomial fit was then calculated from these fixations and applied online to adjust eye position to precisely align the scotoma to the participant’s fovea. This procedure has been described in detail elsewhere (Laurin et al., 2025; Léné et al., 2020; Song et al., 2021).

The scotoma was shaped like a circle with a radius of 2.5° with a circular notch (1.5° radius) removed, extending 1° from the scotoma center to the edge of the notch, oriented at −135° from the rightward horizontal (Figure 1C). This ‘bitten cookie’ shape was designed to encourage PRL development within the notch (or ‘bite’), where peripheral acuity is best.

**Figure 1.**
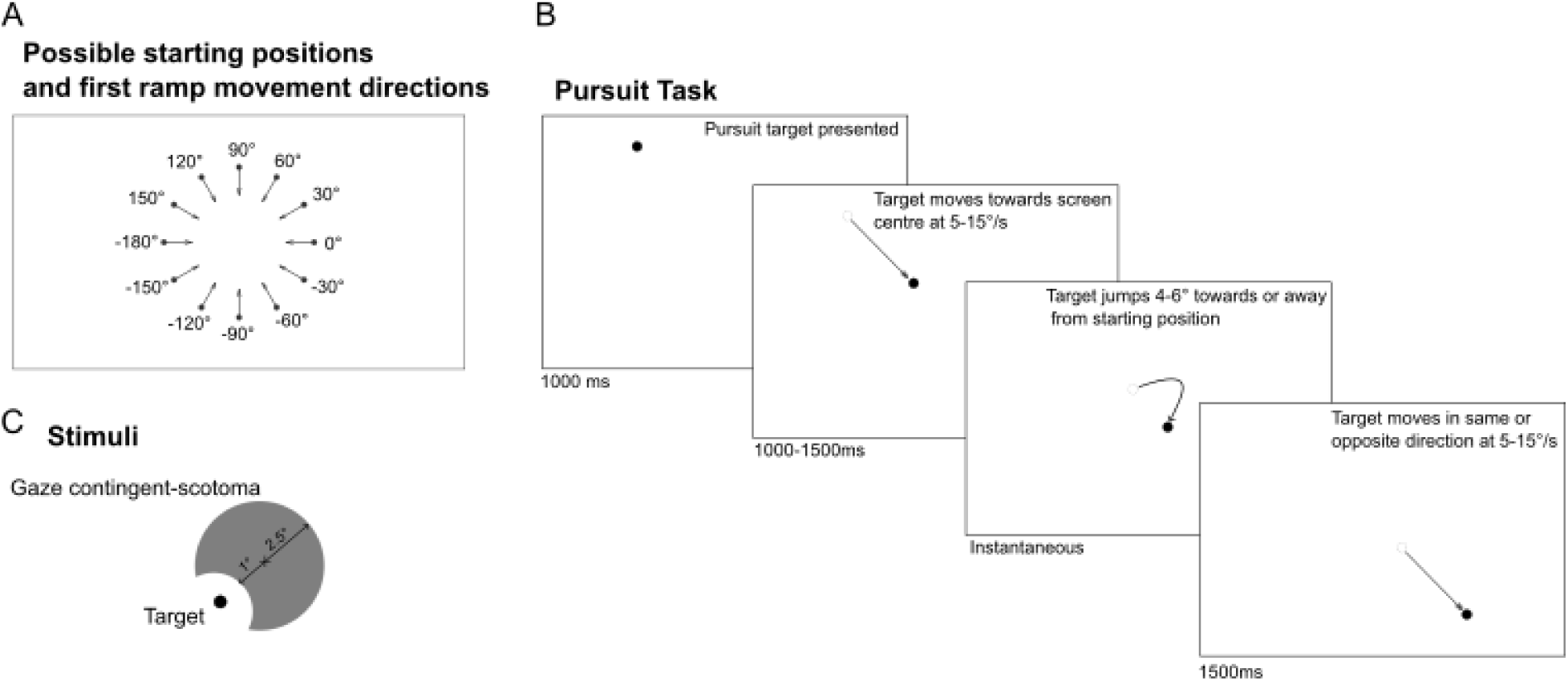
Experimental paradigm. A) All possible starting positions and corresponding first ramp motion directions. Starting positions were spaced 30° apart, at an 8° eccentricity from the center of the screen. B) Pursuit task trail sequence: trials began with a 0.5° black target appearing in one of 12 positions for 1000ms. The target then moved toward screen center at a randomly selected speed (5-15°/s) for a variable duration (1000-1500ms, first ramp), then stepped 4-6° in the same or opposite direction. The target then moved in the same or the opposite direction of the first ramp for 1500ms at randomly selected speed (5-10°/s). C) Scotoma and target parameters. The scotoma has a radius of 2.5°, with a circular notch of 1.5° radius as the ‘bite’ area, oriented −135° from the rightward horizontal. The target radius is 0.25°.

### 2.3 Apparatus and Procedure

We used a modified step-ramp pursuit task (Rashbass, 1961) implemented in Matlab (MathWorks, Inc), with the Psychophysics Toolbox (Brainard, 1997). The control and experimental conditions were identical except for the presence of a gaze-contingent scotoma in the experimental condition. Participants performed multiple blocks of 30 trials each. Each trial began with the appearance of a 0.5° circular black target for 1000ms at one of 12 possible starting positions, evenly spaced at 30° intervals around an 8° radius circle relative to the screen center (Figure 1A). The target then jumped outward from its initial position along the motion axis and moved toward center, crossing the original point 200ms after movement onset (Rashbass, 1961). This step-ramp design is intended to elicit smooth pursuit, rather than a catch-up saccade, as the initial eye movement. Target speed was randomly selected (5-15°/s) and motion continued for a variable duration (1000-1500ms, first ramp), then stepped 4-6° in the same or opposite direction. It then resumed motion for 1500ms at a new randomly selected speed (5-15°/s, second ramp), either continuing in the same direction or reversing (Figure 1B). The velocity difference between the first and second ramp ranged from 0-10°/s. The inter-trial interval lasted 750ms. Participants repeated this task over the course of 5 days across numerous blocks. Day 1 consisted of 5 blocks of the control condition (no scotoma), and 10 blocks of the experimental condition (with scotoma). Days 2-5 each consisted of 10 blocks of the experimental task, followed by a single control condition block on the final day.

### 2.4 Data Pre-processing

We collected a total of 4,500 control trials and 37,500 experimental trials from the 25 participants. We visually inspected all trials using a custom analysis interface in Matlab. Trials were excluded if the participant failed to track the target (e.g., due to distraction), if a blink occurred between 100ms before the target step and the first catch-up saccade following the step, if the catch-up saccade had more than one velocity peak, if a saccade occurred during the target step, if the saccade was in the opposite direction to target direction, or if eye recording failed. This left us with 26,050 experimental trials (69%) and 3,268 control trials (73%).

Saccade latency was calculated as the time between the target step and the onset of the first following catch-up saccade. For saccade latency analyses, we excluded trials with catch-up saccades that occurred less than 90ms after the target step as we considered them to be responses to the preceding ramp rather than to the step itself (de Brouwer et al., 2002). This left 20,565 experimental trials (55%) and 2,557 control trials (58%). However, for saccade amplitude and endpoint error analyses, we included trials with the first saccade occurring less than 90ms after the step if they included a secondary saccade as these would be directed to the target after the step, given that the amplitude of the catch-up saccade is determined by the sensory error 100ms before the saccade rather than by the step as is the case for latency (de Brouwer et al., 2002). In these cases, we did not analyze the first saccade but instead analyzed the second saccade. To be clear, we analyzed only the second saccade in these trials if the first saccade to the step had a latency of less than 90ms. If the latency of the first saccade was 90ms or greater, we used only the first saccade. This allowed us to preserve data from trials that were discarded based on the latency analysis. Therefore, in each trial included for these analyses, the endpoint error and amplitude of a singular catch-up saccade was analyzed, either the first or second directed to the step. The saccade selected was conditional on the latency of the first saccade to the step.

Eye position was low-pass filtered using an autoregressive forward-backward filter with a cut-off frequency of 50 Hz. Eye velocity and acceleration were derived from position signals using a central difference algorithm (±10ms window), and saccades were detected using an acceleration threshold of 500 deg/s^2^. The catch-up saccade analyzed in each trial was the first saccade directed to the target step after the first ramp of motion. Among the measures analyzed for these catch-up saccades were amplitude and peak velocity. Because the commands for saccades and pursuit add together during catch-up saccades, saccade amplitudes and peak eye velocities must be corrected to isolate the saccade component and remove the smooth component (de Brouwer et al., 2002). More specifically, saccades made in the same (forward saccades) or opposite (reverse saccades) direction of the smooth eye movement have differing amplitudes, but do not differ once the smooth component is removed. This is done by subtracting pursuit eye velocity integrated over the saccade duration from saccade amplitude, leaving us with saccade amplitude computed purely from the saccadic command. Saccade amplitude (S_AMP_) was first computed as the Euclidian distance between eye position at saccade onset (*EYE_ONX_*, *EYE_ONY_*) and offset (*EYE_OFFX_*, *EYE_OFFY_*)

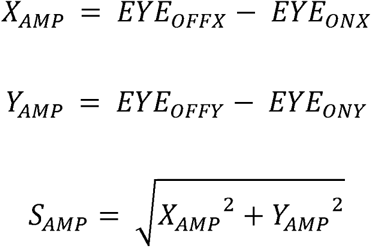

The smooth component of the saccade amplitude was then removed by subtracting the product of the saccade duration (S_DUR_) and mean pursuit velocity (V_PURS_) from this value (de Brouwer et al., 2002). Saccade duration (S_DUR_) was the time between saccade onset and offset. To compute the mean pursuit velocity during the saccade (V_PURS_), we calculated the average pursuit velocity at the ranges 75 to 25ms before saccade onset (V_ON_) and 25 to 75ms after saccade onset (V_OFF_). Averaging over 50ms windows was done to allow for a more accurate estimate of velocity signals from noisy data (de Brouwer et al., 2002).

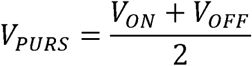

Saccade peak velocity was computed by subtracting the pursuit velocity (V_PURS_) from the maximum velocity during the saccade (V_MAX_).

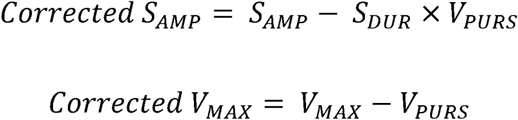

We also measured saccade endpoint errors (S_ERR_), which were calculated as the Euclidian distance between the eye landing position and the target at the time of saccade offset. We computed both horizontal and vertical endpoint errors (ERR_X_, ERR_Y_) from the horizontal and vertical eye (EYE_OFFX_, EYE_OFFY_) and target positions (TAR_OFFX_, TAR_OFFY_).

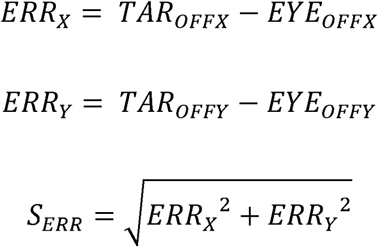

To quantify the degree of alignment of saccade endpoint error and amplitude vectors with the bite direction, each endpoint error (S_ERR_) and corrected amplitude vector (Represented by the 2-D displacement vector of the saccade with a magnitude of the corrected amplitude: Corrected S_AMP_) were projected onto the axis defined by the bite orientation (*θ_BITE_*: −135° from rightward horizontal). Angular direction vectors for endpoint error (*θ_S_ERR__*) and amplitude (*θ_S_AMP__*) were obtained using the horizontal and vertical components used to calculate each variable. For endpoint errors, this projection (Bite axis endpoint error: *ERR_BITE_*) yields a signed scalar for each saccade: positive values indicate a shift from the center of the scotoma toward the scotoma edge in the bite direction, and negative values indicate a shift away from it into the scotoma center, opposite from the bite location.

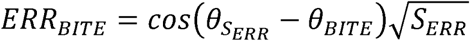

Figure 2 demonstrates the use of this method for computing the bite axis endpoint error from a raw saccade endpoint error between the target and the fovea. Corrected saccade amplitude was similarly projected onto the bite axis (*AMP_BITE_*) using the same method.

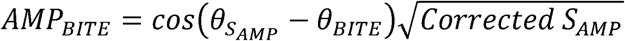

**Figure 2.**
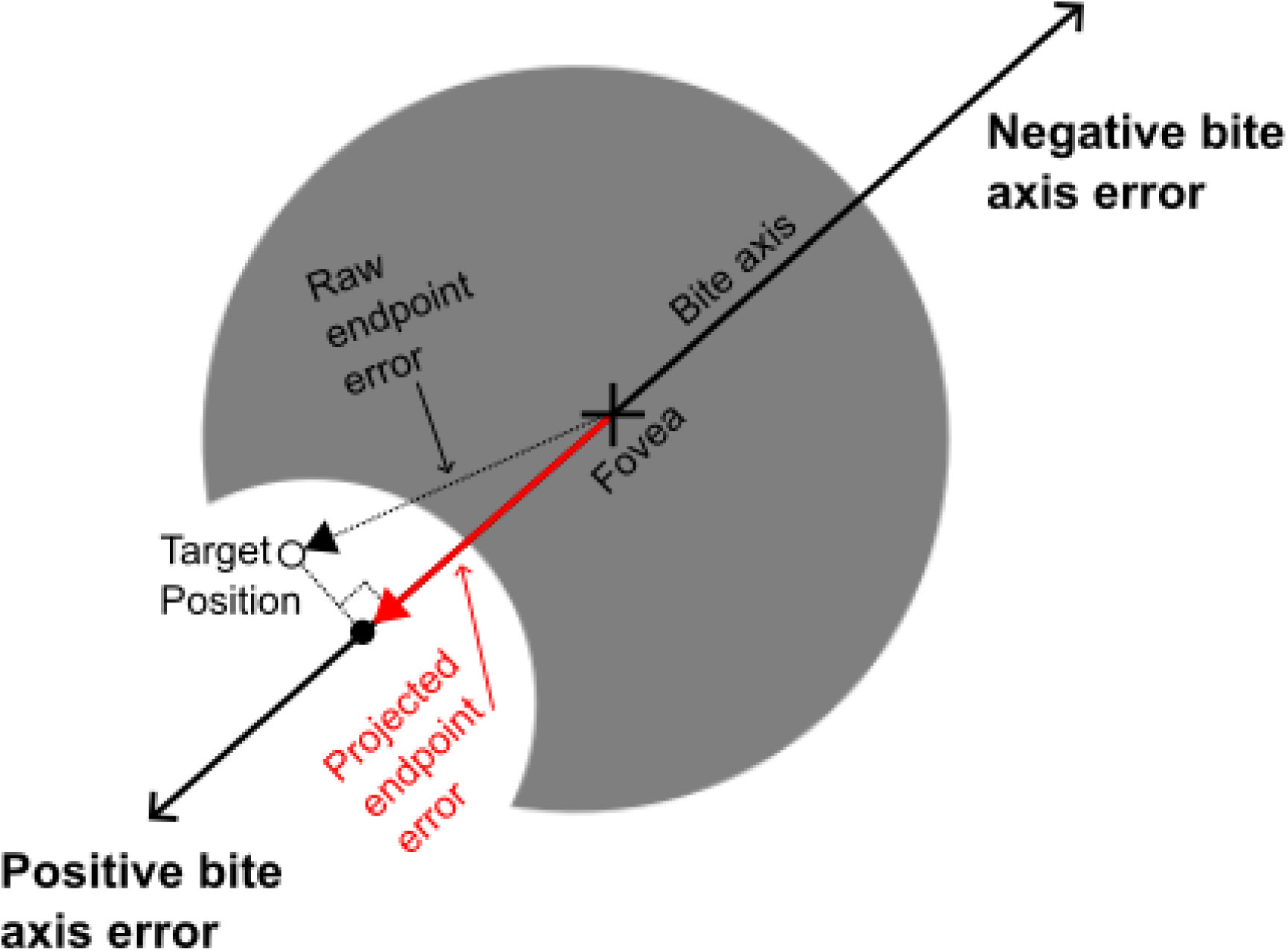
Endpoint error projection onto the scotoma bite axis. The fovea and scotoma positions are shown, along with an example saccade endpoint error vector (target position minus actual endpoint). This raw error vector is projected onto the scotoma bite axis to yield a signed bite axis endpoint error: positive values indicate an error directed toward the bite, and negative values indicate an error directed away from the bite. With adaptation, errors are expected to become progressively more positive; without adaptation, error should be roughly equally positive and negative. This projection collapses the two-dimensional endpoint error into a single value along the bite axis, providing a measure of the degree of saccadic adaptation, or re-referencing, toward the scotoma.

### 2.5 Statistical Analysis

We conducted paired t-tests to compare differences in bite axis endpoint errors and deviation along the PRL axis between training days 1 and 5 within participants. For each participant, the mean bite axis endpoint error was also calculated separately for each of the four target direction quadrants on each of the five training days. Target directions were divided into four quadrants. We conducted a two-way repeated-measures analysis of variance (ANOVA) to assess the effects of Day (Day 1 vs. Day 5) and Quadrant (Q1–Q4) on bite axis endpoint error across all participants. Post-hoc paired comparisons were corrected for multiple comparisons using the Holm–Bonferroni procedure. Day 1 versus Day 5 comparisons were conducted separately for each of the four quadrants. Pairwise comparisons between the four quadrants were performed after collapsing days within each quadrant. Holm–Bonferroni correction was applied separately to each family of comparisons.

For each participant, we ran regressions on bite axis saccade amplitude, using the bite axis retinal slip and position errors as the two independent variables. A separate regression was run for each quadrant (either Q1 or Q3) on all days for each participant. We also compared across days within quadrants by running regressions separately for quadrant 1 and 3 on both days 1 and 5. Paired t-tests were used to compare regression coefficients between quadrants and days.

We conducted a repeated measures ANOVA comparing bite axis endpoint errors for pre-saccadic visibility conditions across day, as well as Holm-Bonferroni corrected posthoc paired t-tests between day 1 and 5.

A repeated-measured ANOVA was also conducted to compare saccade latencies for different visibility conditions across training days. Holm-Bonferroni corrected posthoc paired t-tests between day 1 and 5 followed for each of these visibility groups.

Finally, we ran a repeated measures ANOVA to compare the root mean squared deviation of the linear fit of each individual’s amplitude and peak velocity main sequence on Day 1, 5 and the control session. Paired t-tests between each day followed.

## 3. Results

The primary aim of our study was to examine whether, following training with an artificial central scotoma, catch-up saccade endpoints shift to reflect the use of the PRL in place of the fovea as the oculomotor reference. During training, we observed that catch-up saccades landed in such a way that either the target was occluded by the scotoma, was visible outside of the scotoma borders, or it was visible in the PRL (bite area). Figure 3 depicts eye position traces from a typical trial from the middle of training (day 3). In panel A are the horizontal and vertical eye positions over time, with saccades highlighted in bold overlaid on the pursuit traces, while the velocity is plotted in panel B. In panel C, the spatial position of the eye and target are plotted, with markers for eye position sampled every 10ms to visualize the saccade versus pursuit components of the trial.

**Figure 3.**
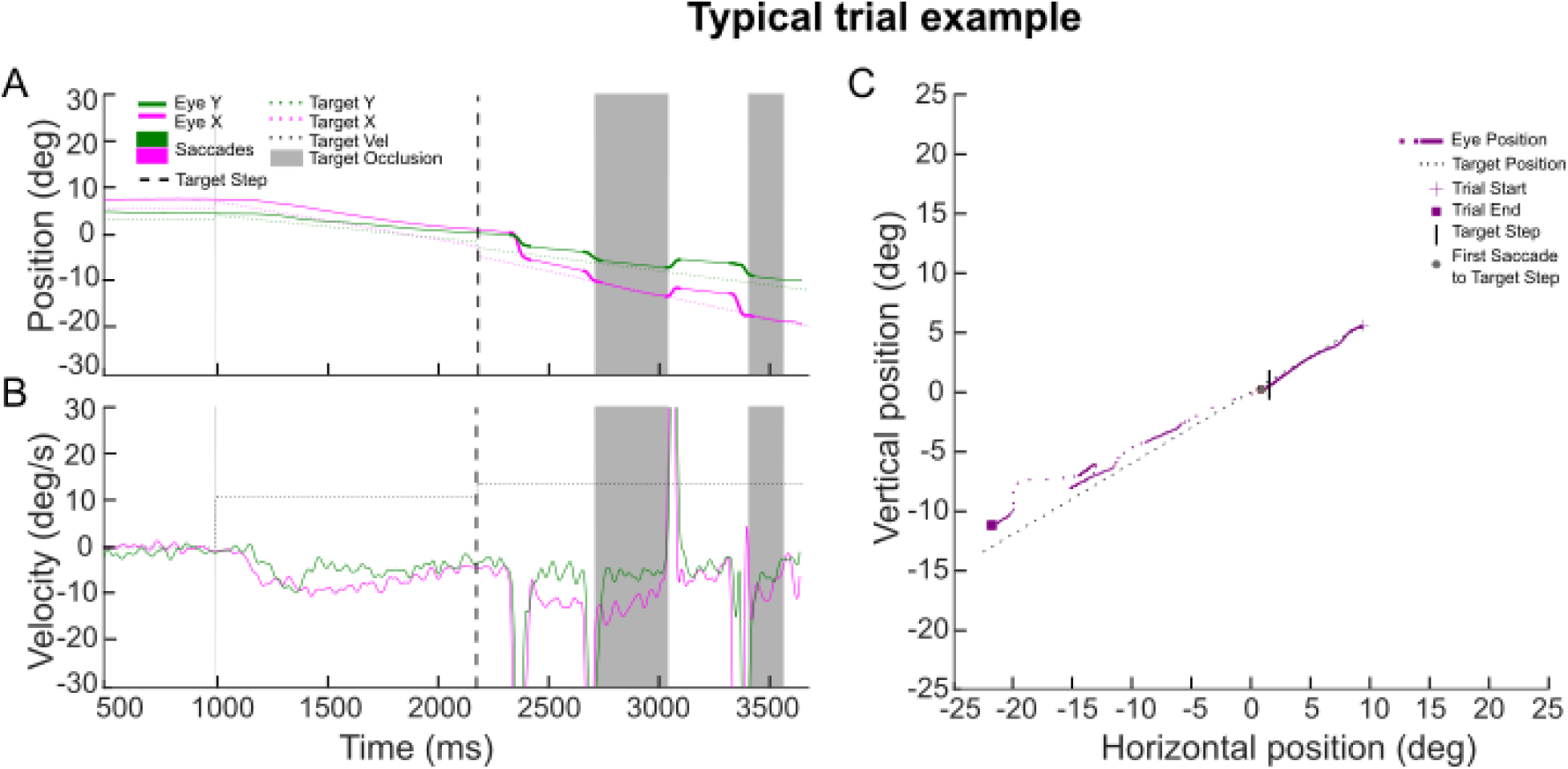
Typical trial. Example of a typical trial for one participant on day 3 of the experiment. In panels A and B, the horizontal (green) and vertical (pink) components of eye position, target position, eye velocity and target velocity (black dotted) are plotted against time. The black vertical dashed line is the time of the target step. Following the target step, there are four highlighted bold portions of each line in panel A, indicating saccades. Gray shaded areas depict the time during which the target was occluded behind the scotoma. In panel C, the eye and target spatial positions are plotted in purple and black, respectively. To facilitate visualization of movement dynamics, eye position traces (purple) are plotted as one sample every 10ms, enabling clearer distinction between smooth pursuit and saccadic displacements. The fixation cross depicts the start of target motion during the trial, the black vertical bar indicates the time of the beginning of the target step, th grey circle indicates the beginning of the first catch-up saccade to the step, and the purple square represents the end of the trial.

In this trial, the first saccade after the target step landed such that the target was visible at the saccade endpoint. Another catch-up saccade followed landing so that the target became occluded (gray shaded region), requiring another corrective third saccade to regain target visibility. The target remained visible at the onset of the fourth saccade but then was occluded again when it landed.

### 3.1 Saccade endpoints shift towards scotoma bite over training days

Figure 4 displays the catch-up saccade endpoint errors (i.e. the saccade landing position with respect to the target position) during each trial for the first saccade for day 1 (red) and day 5 (blue), with a depiction of the scotoma underlaid for reference, for three different participants.

**Figure 4.**
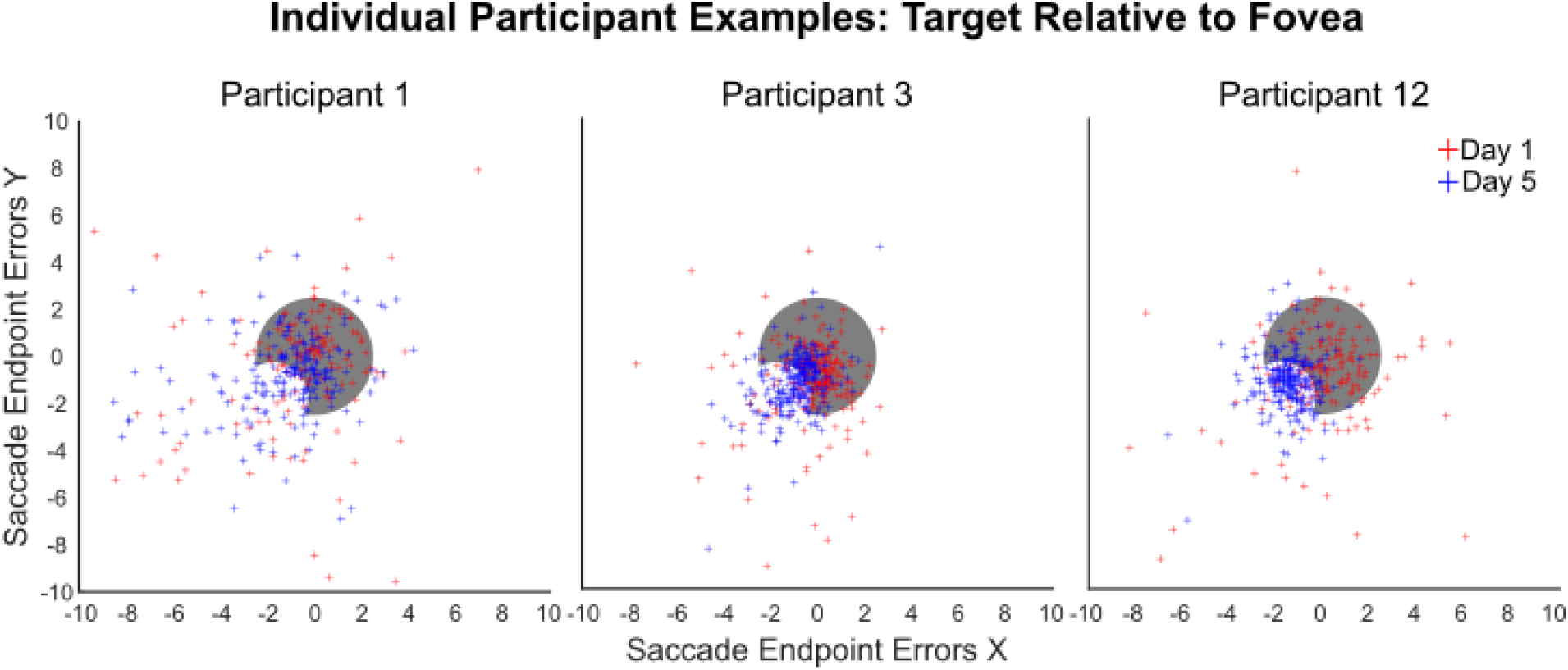
Three individual participant endpoint errors on day 1 and 5 of training. Horizontal and vertical endpoint errors for the catch-up saccade plotted on top of an image of the scotoma used in the experiment. Only the trials included in any saccade endpoint error related analyses are included in these visualizations.

**Figure 5.**
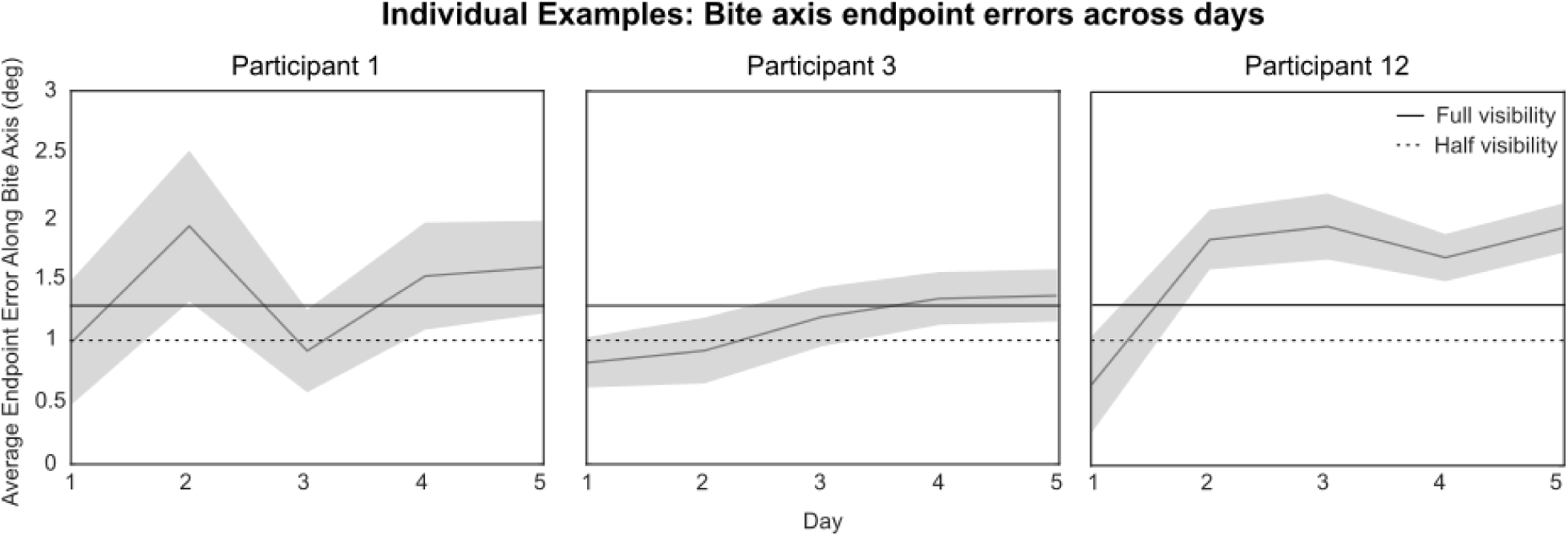
Individual participant bite axis endpoint errors over training days. Grey lines denote the average bite axis endpoint error for each day for 3 participants, with a light grey shaded 95% confidence interval. The solid black horizontal line marks the approximate threshold for full target visibility (the target lies entirely outside the scotoma), while the dotted black line marks the approximate threshold at which half of the target is visible and the other half remains occluded.As can be seen, learning curves varied between participants. Some participants learned very quickly, while others were more gradual.

Horizontal and vertical saccade endpoint errors of 0 indicate that the saccade brought the target exactly onto the fovea, which was occluded by the scotoma. Endpoint errors closer to the origin (0) were thus expected when the fovea was used as the oculomotor reference. The target was visible only when endpoint errors either exceeded the scotoma radius in both horizontal and vertical directions or when errors placed the target within the PRL (bite area) in the lower left quadrant. In this case, the target was visible when the endpoint errors exceeded the smaller bite radius rather than the full scotoma radius. On the first day of training (red crosses), participants made more saccades to align the target to the fovea, and so the target was occluded by the scotoma. Participants also made saccades so that the target was visible outside of the scotoma but towards locations other than the bite/PRL. By the last day of training (blue crosses), some participants, such as participant 12, showed clustering in the bite location, indicating a shift in saccade endpoints from aligning the target to the fovea to aligning it to the PRL. Between the first and last day of training, we observed a shift over the course of training days for some participants in the direction of the scotoma bite. However, there was variability in performance between participants.

If adaptation occurred, saccade endpoint errors were expected to shift in the direction of the scotoma bite over the five days of training. To quantify this shift, we projected the endpoint errors onto the axis that the bite was oriented (Figure 2). This allowed us to quantify the degree to which endpoint errors were aligned with the scotoma bites axis of orientation. Positive projected bite axis endpoint errors represented endpoint error locations that were more aligned with the direction where the bite was located while negative values were associated with the opposite direction, away from the bite. With adaptation, errors are expected to become progressively more positive; without adaptation, errors should be roughly equally positive and negative. Figure 5 shows the projected endpoint errors as a function of training day for the same three participants as in Figure 4. Here we see an increase in bite axis endpoint errors between the first and last day or training for each participant, however with different learning curves.

Paired t-tests indicated that 13/25 participants showed a significant difference in bite axis endpoint errors between days 1 and 5 of the experiment (mean = 0.50°, min = -0.30°, max = 1.54°). This data is displayed in table 3-1.

We observed similar results when examining the bite axis endpoint errors across days for all participants combined (Figure 6). A paired t-test showed a significant difference in bite axis endpoint errors between day 1 and day 5, (t(24) = −5.657, p < 0.0001, day 1 mean = 0.53 deg, day 5 mean = 1.03 deg, mean difference = 0.50 deg). Average bite axis endpoint errors exceeding 1° indicate that for most trials, at least half of the target is visible at the saccade endpoint, while errors exceeding 1.25° indicate that the target is fully visible. As can be seen in figure 6 and table 1, although some participants did not show a sigificant difference in bite axis endpoint errors between day 1 and 5, most participants (16/25) maintained at least half target visibility on the final day of the experiment.

**Figure 6.**
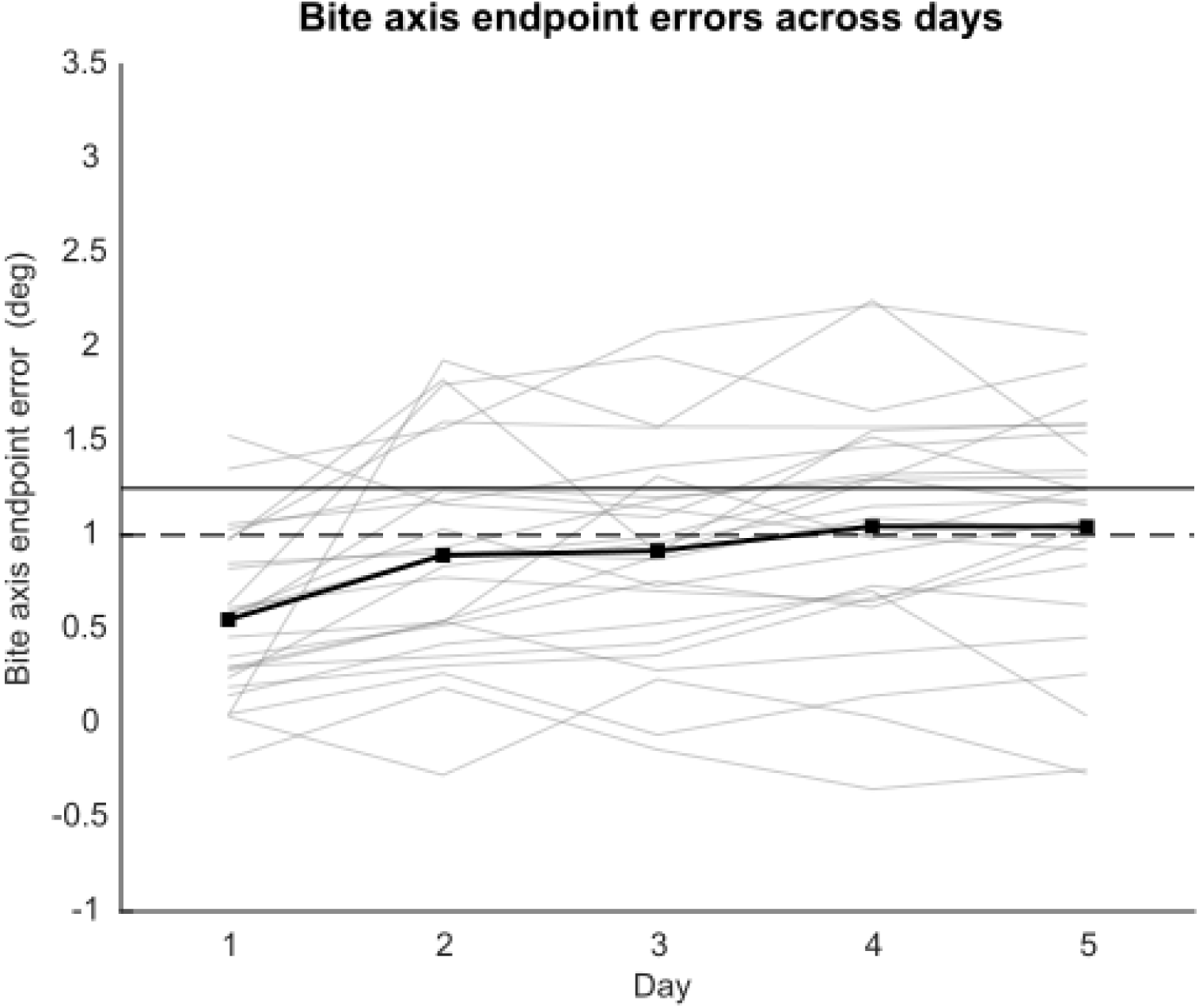
Average participant bite axis endpoint errors over training days. Filled squares show the mean endpoint error for each day. Grey lines show individual endpoint curves for each participant. The dashed horizontal line is the approximate threshold for a bite axis endpoint error shift of 1°, which would result in half the target being occluded by the scotoma. The filled horizontal line at 1.25° is the approximate threshold for full target visibility.

**Table 1.** Individual participant average bite axis endpoint errors and paired t test results. 13/25 participant showed statistically significant increases in bite axis endpoint errors from the first to last day of training.

| Participant | Day 1 | Day 5 | P value | Shift |
| --- | --- | --- | --- | --- |
| 1 | 0.98 | 1.59 | 0.102 | 0.61 |
| 2 | 0.30 | 1.74 | 0.000*** | 1.44 |
| 3 | 0.82 | 1.36 | 0.001** | 0.53 |
| 4 | 0.05 | 1.46 | 0.000*** | 1.41 |
| 5 | 0.32 | 0.65 | 0.217 | 0.32 |

*Table 1. Individual participant average bite axis endpoint errors and paired t test results.*
|  |  |  |  |  |
| --- | --- | --- | --- | --- |
| 6 | 1.03 | 1.47 | 0.018* | 0.44 |
| 7 | 0.28 | 0.43 | 0.386 | 0.15 |
| 8 | 0.46 | 1.24 | 0.001** | 0.79 |
| 9 | 0.59 | 1.07 | 0.021** | 0.48 |
| 10 | 0.58 | 1.05 | 0.098 | 0.47 |
| 11 | 1.03 | 1.30 | 0.211 | 0.27 |
| 12 | 0.65 | 1.91 | 0.000*** | 1.26 |
| 13 | 0.16 | 0.02 | 0.646 | -0.14 |
| 14 | 0.38 | 0.99 | 0.001** | 0.60 |
| 16 | 0.19 | 0.81 | 0.003** | 0.62 |
| 16 | 0.03 | -0.26 | 0.119 | -0.29 |
| 17 | -0.30 | -0.06 | 0.043** | 0.24 |
| 18 | 0.05 | 0.26 | 0.204 | 0.21 |
| 19 | 1.45 | 1.25 | 0.490 | -0.21 |
| 20 | 0.54 | 0.94 | 0.160 | 0.40 |
| 21 | 0.24 | 1.17 | 0.000*** | 0.92 |
| 22 | 0.73 | 1.21 | 0.333 | 0.47 |
| 23 | 0.89 | 1.03 | 0.747 | 0.15 |
| 24 | 1.28 | 2.10 | 0.003** | 0.81 |
| 25 | 0.56 | 1.17 | 0.002** | 0.61 |

### 3.2 Directional asymmetries in catch-up saccade adaptation

Next, we examined differences in bite axis endpoint errors depending on pursuit direction of the first ramp of motion. Given the shape of the scotoma, we predicted that if adaptation occurred, bite axis endpoint errors would differ depending on whether the target moved toward, away from or orthogonal to the bite. As described in methods and figure 2, positive bite axis endpoint errors are indicative of adaptation towards the PRL. Specifically, we expected endpoint errors to be largest when the target moved into bite and scotoma (towards cartesian quadrant 1), and smallest away from the bite (towards cartesian quadrant 3). This is because saccades made to motion directed into the bite would require an overshoot for the target to land in the bite than saccades made to motion directed away from the bite, which would require a slight undershoot. Trials were grouped into pursuit direction quadrants based on the first ramp of motion. Figure 7A displays the average bite axis endpoint errors for each pursuit direction quadrant for all participants on each day of training, with the standard deviation included in the error bars. A two way repeated measured ANOVA with training day (first versus last) and pursuit direction quadrant on bite axis endpoint error revealed a significant main effect between training day (F(1, 24) = 34.85, p < 0.001) and quadrants (F(3, 72) = 24.70, p < 0.001), but not the interaction (F(3, 72) = 0.85, p = 0.47). Holm-Bonferroni corrected pairwise comparisons showed that endpoint error increased significantly from day 1 to day 5 in all four quadrants (Q1: t(24) = -3.96, p = 0.002; Q2: t(24) = −5.89, p < 0.001; Q3: t(24) = -2.73, p = 0.012; Q4: t(24) = -3.17, p = 0.008).

**Figure 7.**
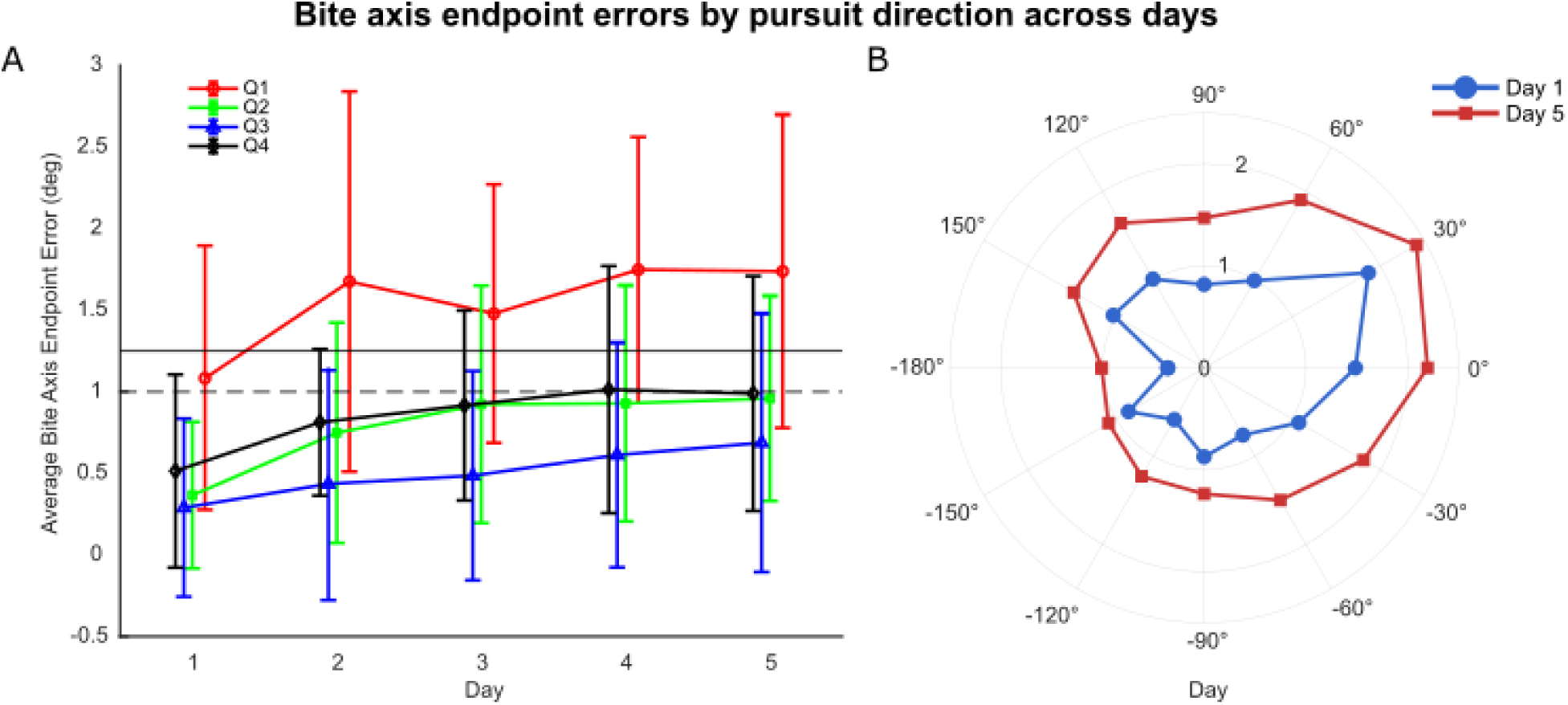
Average participant bite axis endpoint errors grouped by pursuit direction over training days with standard deviation error bars. A) Pursuit direction quadrants are defined relative to horizontal: Quadrant 1 (0–90°) is toward the top right of the screen, opposite the scotoma bite; Quadrant 2 (90–180°) is toward the top left, orthogonal to the bite; Quadrant 3 (180–270°) is toward the bottom left, movement is away from the bite; and Quadrant 4 (270–360°) is toward the bottom right, orthogonal to the bite. The dashed horizontal line marks the threshold for at least 50% target visibility, and the solid horizontal line marks the threshold for full target visibility. B) Bite axis endpoint errors for every pursuit target direction on Day 1 and Day 5.

Collapsed across days, quadrant comparisons showed that Q1 endpoint errors were significantly higher than all other quadrants (all p < 0.001), while Q3 and Q4 also differed significantly from each other (p = 0.040); Q2 vs. Q3 (p = 0.134) and Q2 vs. Q4 (p = 0.248) were not significant after correction. Bite axis endpoint errors were thus largest with pursuit direction towards quadrant 1, into the bite. However, the increase in bite axis endpoint error from day 1 to 5 still occurred in all pursuit directions used in our task, although with varying magnitudes (Figure 7B). The scotoma shape interacts with the pursuit direction such that saccades made to motion directed into the bite require a larger amplitude for the target to land in the bite than those directed away from the bite. For example, if the pre saccadic position error is 5 degrees for a trial where the target moved towards quadrant 1, the corresponding saccade amplitude would be around 6.5 degrees for the target to land in the bite. A trial with the same pre-saccadic position error with target motion direction towards quadrant 3 would require a saccade of 3.5 degree amplitude to have the target land in the bite. Therefore, saccades directed towards quadrant 1 require an overshoot to the target while those directed towards quadrant 3 require an undershoot. Since catch-up saccades naturally undershoot targets (de Brouwer et al., 2002), this undershoot bias works against the amplitude needed to capture the bite for quadrant 3 motion, though compensation still occurs to some degree. For motion towards quadrant 1, by contrast, the amplitude needed to capture the bite is in the same direction as an overshoot, so saccades in this direction more readily exceed target position. This directional overshoot for quadrant 1 may also larger but less accurate saccades, which could explain the larger bite axis endpoint errors observed in that direction compared to others.

### 3.3 Saccade amplitude changes: Pursuit direction along bite axis

To further quantify saccadic adaptation along the bite axis, we isolated trials with pursuit direction toward quadrant 1 or quadrant 3, since these directions align most closely with the bite axis itself and therefore provide the clearest signal for adaptation along this dimension. This approach allowed us to assess whether the saccadic system was appropriately scaling its motor output to the pre-saccadic retinal error along the bite axis, and whether this relationship changed with training, which would provide a more direct, trial-by-trial measure of re-referencing than the day-level comparisons alone.

We projected the 2-D saccade displacement vector onto the bite axis, with the saccade amplitude corrected for the smooth component used as the magnitude. We also projected the 2-D position error and retinal slip (velocity error) vectors centered 100ms before the saccade along the bite axis as well. For each participant, we ran regressions on this bite axis saccade amplitude, using the bite axis retinal slip and position errors as the two independent variables. The regression equation is as follows:

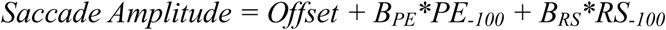

A separate regression was run for each pursuit direction quadrant (either Q1 or Q3) on all days for each participant. As described in the previous section and shown in figure 7, the largest difference in bite axis endpoint errors by pursuit direction occurred between opposite directions along the bite axis; errors were largest for pursuit toward quadrant 1 (into the scotoma bite) and the smallest for pursuit toward quadrant 3 (away from the scotoma and bite). Trials in these opposing directions were therefore analyzed separately. Figure 8 displays, for a single participant, the projected position error sampled 100ms before the catch-up saccade as a function of its corrected projected amplitude, with their respective regression lines for each direction overlayed. The data and regression line for pursuit direction towards quadrant 3 intercepts roughly at 0, while quadrant 1 direction is offset in the negative direction by several degrees. This offset reflects the larger endpoint errors observed for trials moving into the scotoma bite (towards Q1), consistent with the need to overshoot the target in that direction. The regression slopes were similar for both directions, meaning saccade amplitude scaled with position error in similar ways regardless of pursuit direction. The offset of the regression line between quadrant 1 and 3 however captures the directional asymmetries, consistent with results from the previous section.

**Figure 8.**
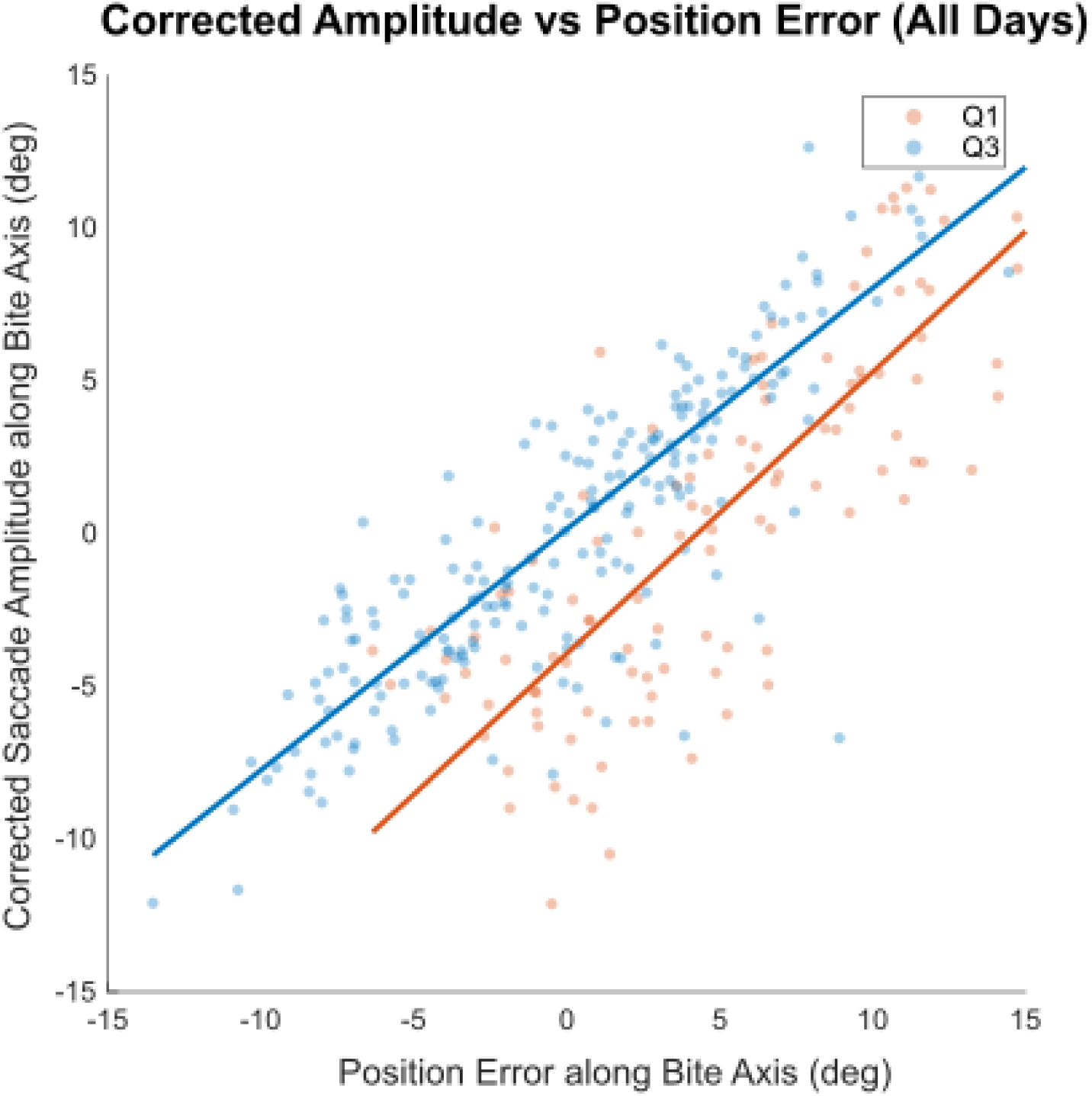
Corrected Amplitude vs. Position error example. Single participant example of position error 100ms before saccade (projected onto the bite axis) as a function of corrected saccade amplitude (projected onto the bite axis) for quadrant 1 and 3 pursuit directions on all training days. Regression lines for each direction are overlayed over the scattered data.

Regression coefficients including the data from all days for both quadrants 1 and 3 are compared in panels A-C of Figure 9. Panel A displays the regression intercept for each participant with pursuit directions in quadrants 1 and 3. Panels B and C compare the regression coefficients for the sensory variables included in the regression for each quadrant. In this regression, the slope coefficients (*B_PE_* and *B_RS_*) explain the sensitivity of saccade amplitude to pre-saccadic position error and retinal slip, respectively. Specifically, it reflects how much the saccadic system scales its motor output as a function of sensory errors. The intercept reflects the component of saccade amplitude that is independent of the sensory errors, capturing a constant added bias in the motor output rather than a change in how the oculomotor system uses incoming error signals.

**Figure 9.**
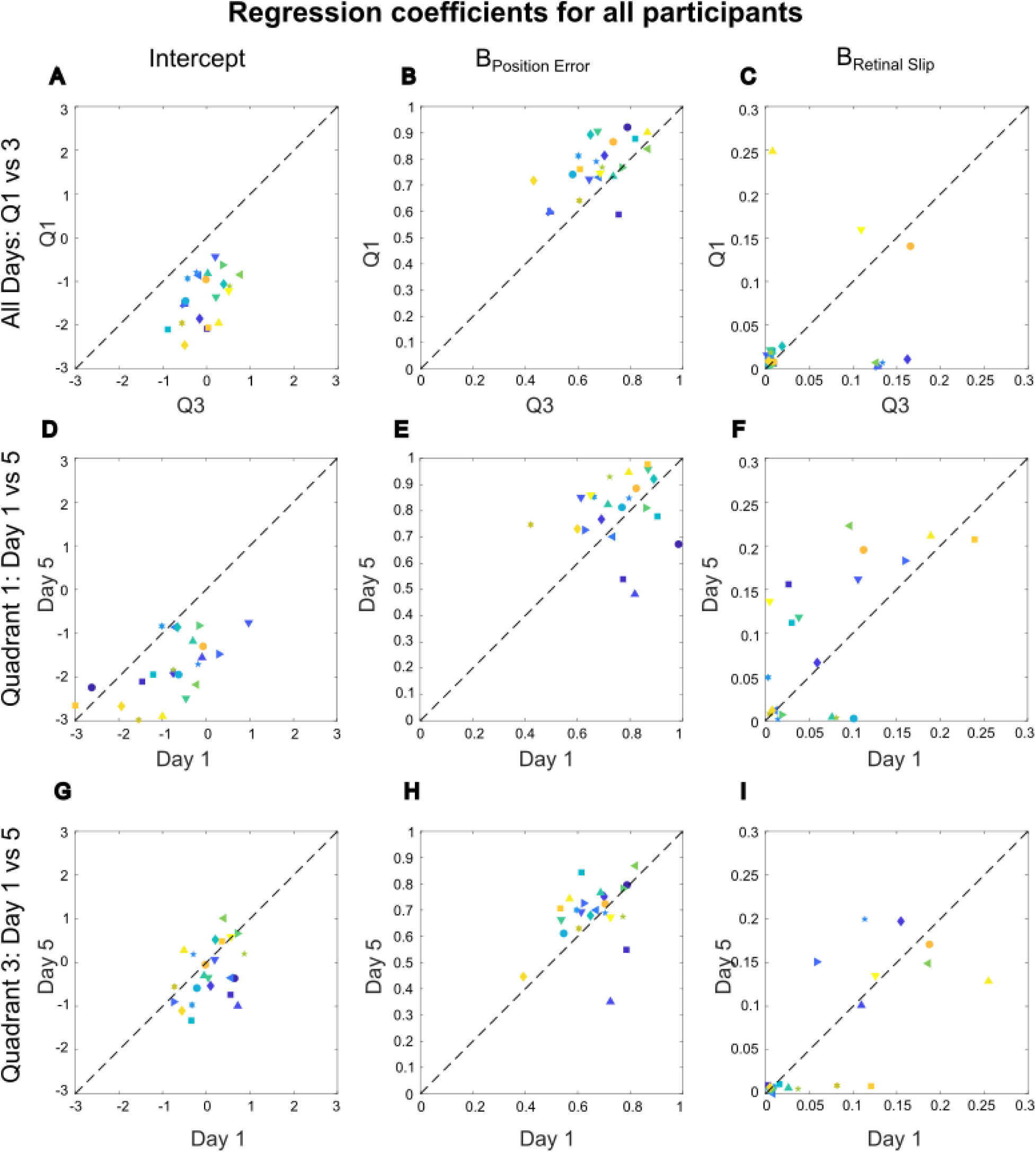
Regression coefficients for pursuit directions along bite axis. Each participant is represented by a unique marker across scatterplots. Panels A-C include data from all days comparing pursuit directions towards quadrant 1 and 3. Panels D-F consist of only quadrant 1 data on day 1 and 5, and panels G-1 only quadrant 3 for days 1 and 5. Position error and retinal slip (velocity error) 100 ms before the saccade was projected onto bite axis as independent variables, and catch-up saccade amplitudes (corrected) projected onto bite axis were the dependent variable for each regression.

To visualize within-participant change in regression coefficients between quadrants, we computed the signed orthogonal distance of each participant’s paired coefficient values from the identity line. We estimated the probability density of these values across participants using Gaussian kernel density estimation separately for each coefficient (intercept, position-error, retinal-slip) (Figure 10A). To allow for comparison of coefficients with different scales within a single plot, each density was rescaled such that zero (indicating no change between conditions) was aligned across coefficients; actual coefficient units are indicated on the corresponding axis beneath the plot in the same colour as their distributions. As can be seen by the difference distribution of regression parameters between days 1 and 5 plotted in figure 10A, there is a very small change in the deviation from the unity line for position error (green distribution), but not retinal slip coefficients (blue distribution) between pursuit directions. However, the deviations for the regression intercepts (red distribution) are much larger, with a shift similar in magnitude to the example in figure 8. This can be interpreted as a shift in saccade amplitude due to overshooting and undershooting along the bite direction, while the influence of retinal slip remains relatively constant between both motion directions. Because the slope relating saccade amplitude to position error was similar across quadrants, this asymmetry cannot be explained by a difference in how strongly the saccadic system responds to a given position error. Instead, the more negative intercept for quadrant 1 indicates that saccades in this direction consistently traveled farther than the position error alone would predict, consistent with a compensatory adjustment that overshoots the target location when pursuit moves into the bite. The near zero intercept for quadrant 3 suggests that little to no such compensation was needed when pursuit moved away from the bite, in line with the smaller endpoint errors observed in that direction. Taken together, this pattern suggests that adaptation along the bite axis is produced primarily as a directionally specific offset in motor output, added on top of a relatively stable relationship between sensory errors and amplitude scaling.

**Figure 10.**
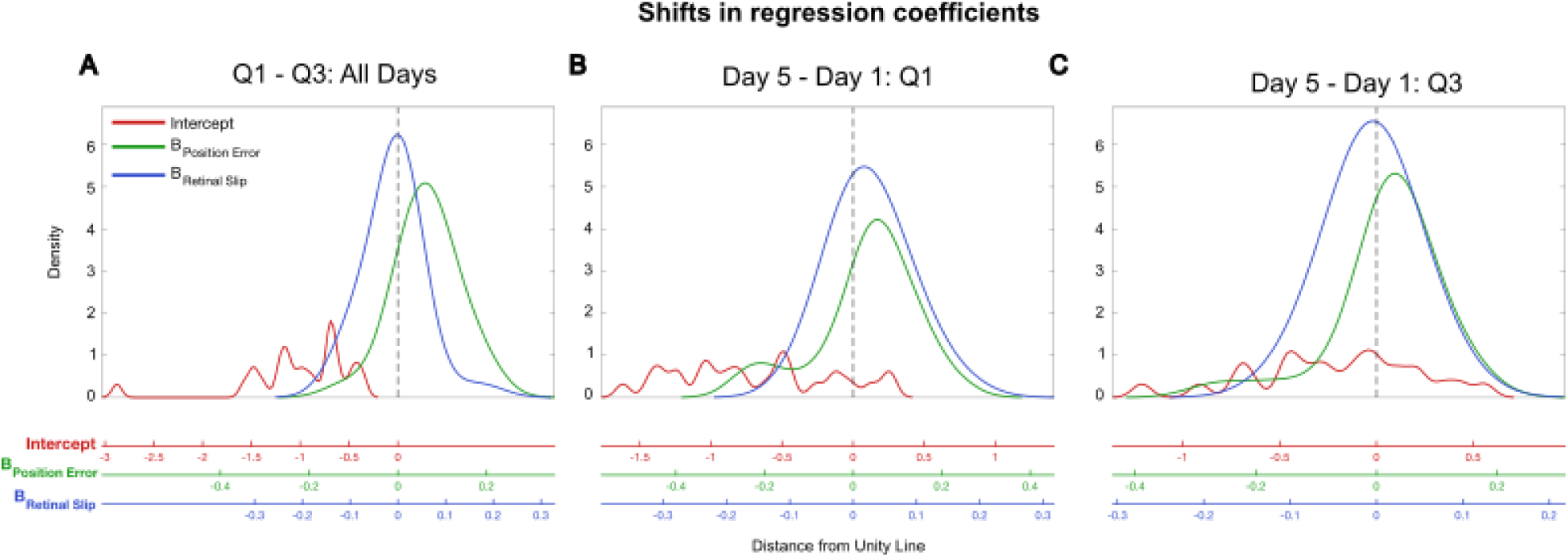
Kernel density estimates of change distributions across conditions for all participants. The distribution of signed orthogonal distances of each participant’s paired coefficient values from figure 9 (intercept, position error and retinal slip) from the identity line was computed comparing quadrants 1 and 3 on both days in panel A, and between days 1 and 5 for both quadrant 1 (panel B) and quadrant 3 (panel C). The probability density of these distributions were computed using Gaussian kernel density estimation (bandwidth = 0.05).

We compared regression coefficients between both quadrants using paired t-tests. The intercepts differed significantly between quadrants (t(24) = -9.352, p < 0.0001), with quadrant 1 showing more negative values. The position error coefficient was significantly higher in quadrant 1 compared to quadrant 3 (t(24) = 4.754, p = 0.0001), suggesting a greater reliance on position error signals with quadrant 1 pursuit direction. The retinal slip coefficients did not differ significantly between quadrants (t(24) = -0.756, p = 0.4569), suggesting comparable use of retinal slip information in both directions.

Additionally, we ran further regressions for each participant using the same dependent and independent variables only using data from a single quadrant, on both days 1 and 5. The scatter plots in the first three columns of figure 9 summarize the results of each regression by plotting the regression fits for each group against each other.

While the first row of figure 9 shows each participants regression coefficients using data from all days, separating quadrant 1 and 3 (Figure 9. A-C), the next rows instead compare coefficients on day 1 and 5 of the experiment separately for quadrant 1 (Figure 9. D-F), and quadrant 3 (Figure 9. G-I), respectively. The distance of each distribution from the unity line with aligned axes to directly compare the group differences across all parameters can also be seen in figure 10, panels B and C.

As expected due to the clear differences in panels A-C of figure 9 and panel A of figure 10, we found larger shift in intercept between day 1 and 5 for trials with target motion towards quadrant 1. We also found significantly different intercepts between day 1 and 5 for motion directed towards quadrant 3, indicating some type of learning over time for both directions along the bite axis (Q1:t(24) = 6.257, p < 0.0001, Q3: t(24) = 2.442, p = 0.0223). There were no differences in position error coefficients between days for either quadrant (Q1: t(24) = −1.396, p = 0.1756, Q3: t(24) = −1.054, p = 0.3025), however there was a small difference in retinal slip coefficients between days for quadrant 1 (t(24) = -2.418, p = 0.0236), but not for quadrant 3 (t(24) = 0.639, p = 0.5287). The most significant difference was between the day 1 and day 5 intercepts for quadrant 1, reflecting both the requirement of overshooting saccades and the robust adaptation over time in that direction, as was seen in figures 7 and 8.

### 3.4 Pursuit Direction Orthogonal to Bite Axis

The previous section quantified amplitude changes for pursuit directions aligned with the bite axis, where adaptation toward the PRL would be expressed as a change in catch-up saccade amplitude along the direction of target motion. However, targets also moved in directions orthogonal to the bite axis, where adaptation cannot be explained by saccade amplitude changes alone. For these movements, adaptation to the PRL would require a change in saccade direction, resulting in an angular shift of the saccade endpoint toward the PRL. Previously, endpoint errors were projected onto a fixed bite-axis vector, which quantified displacement along the scotoma axis but did not isolate directional changes in saccade endpoints relative to the intended target location. Therefore, to further examine whether catch-up saccades adapted toward the PRL for movements orthogonal to the bite axis, we additionally calculated the component of endpoint error oriented along the target-to-PRL vector at the time of saccade landing. Trials in which target motion occurred orthogonal to the bite axis (towards Q2 and Q4) were analyzed, as these directions provide a sensitive measure of angular adaptation because any shift toward the PRL must be expressed as a rotation of the saccade endpoint rather than a change in amplitude. For each trial, a vector was calculated from the target location to the actual saccade endpoint, representing the endpoint error. This vector was projected onto a second vector extending from the target location to the predicted PRL location. The resulting scalar value represents the magnitude of endpoint deviation along the PRL axis, with positive values indicating a shift of the saccade endpoint toward the PRL and negative values indicating a shift away from the PRL.

As shown in Figure 11, median endpoint deviation along the PRL axis increased over training days, with a shift of 0.45° toward the bite axis (group medians: Day 1 = 0.34°, Day 5 = 0.80°). A paired t-test comparing Day 1 and Day 5 median deviations revealed a significant increase in PRL-directed endpoint deviation (t(24) = −5.426, p < 0.001). These findings provide further evidence that catch-up saccades adapted toward the PRL during training, even for target motion directions that were orthogonal to the bite axis and therefore required directional rather than purely amplitude-based adjustments.

**Figure 11.**
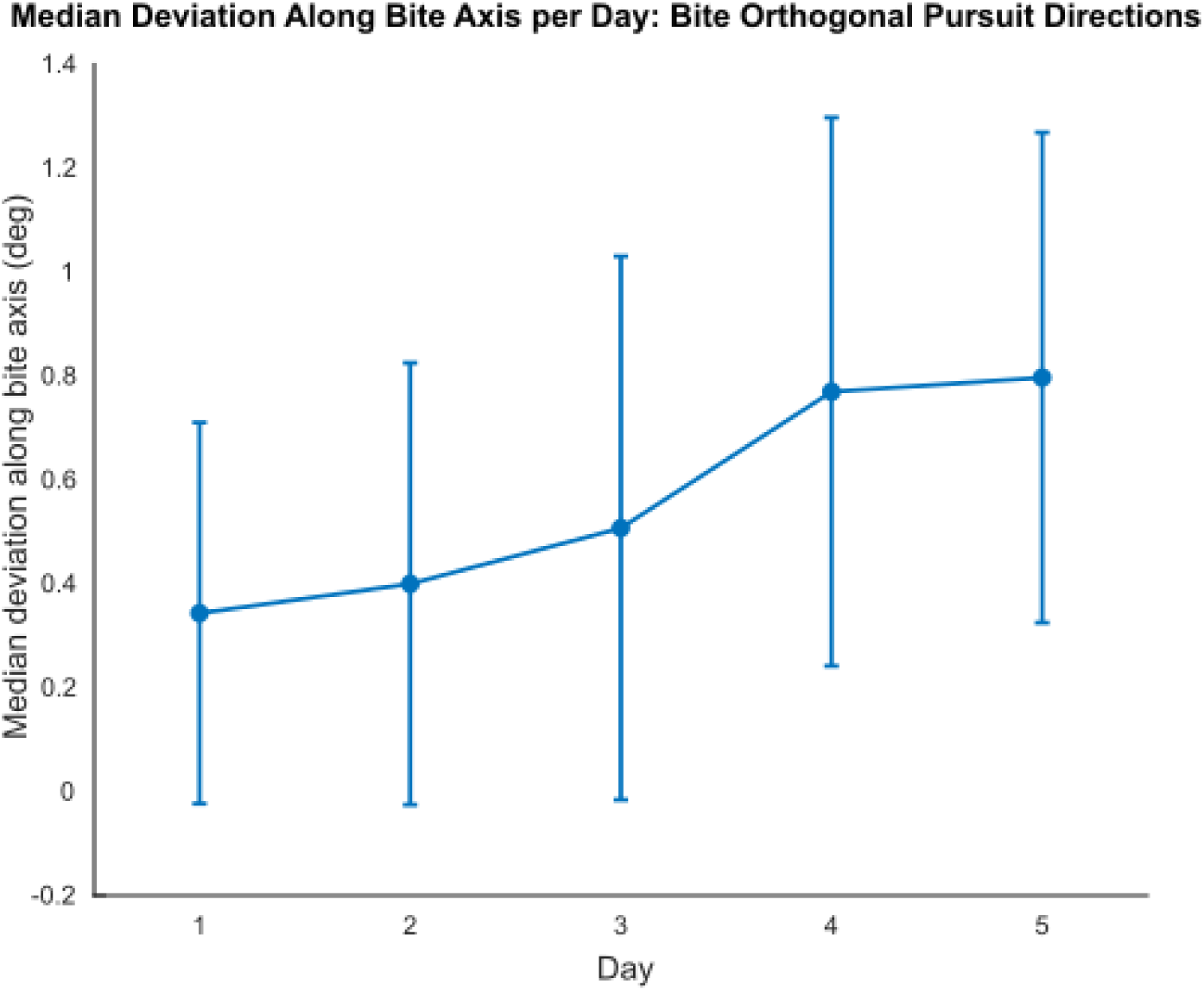
Deviation along bite axis for pursuit direction orthogonal to bite axis. The projection of the fovea-to-saccade-endpoint vector onto the fovea-to-PRL vector wa calculated for each day, providing a measure of saccade-angle deviation along the bite axis. Medians and standard deviations across participants are shown for each day.

### 3.5 Target visibility before the saccade

So far the analysis has focused on how bite axis endpoint errors of catch-up saccades have changed across training days. To fully understand how simulated central vision loss influences catch-up saccade programming and triggering, we also need to consider target visibility during the period in which the saccade is being planned. Visual tracking with a scotoma leads to situations where the target may or may not be visible at critical time points during a task. In our experiment, this critical time point is the target step, which is designed to trigger a catch-up saccade.

We examined whether target visibility during the 90-ms saccade-planning window following the target step influenced saccade landing position relative to the scotoma. This window was selected because it has been shown that the saccadic system requires a minimum of 90 ms to take into account changes in target trajectory (de Brouwer et al., 2002). As shown in figure 12, we grouped trials by whether the target was visible at any point during this window (post-target step) across training days. A repeated measures ANOVA comparing bite axis endpoint errors for pre-saccadic visibility conditions across days revealed a significant effect of day (*F*(1, 24) = 18.177, p<0.0001) but did not reveal a significant main effect of pre-saccadic visibility (*F*(1, 24) = 0.54828, p=0.4662). There was a significant interaction effect between day and pre-saccadic visibility (*F*(4, 96) = 5.3128, p<0.001). Holm-Bonferroni corrected posthoc paired t-tests indicated a sigificant increase in bite axis endpoint errors from day 1 to 5 for both visible (t(24) = -4.851, p = 0.0001) and nonvisible conditions (t(24) = -6.936, p = 0.0000).

**Figure 12.**
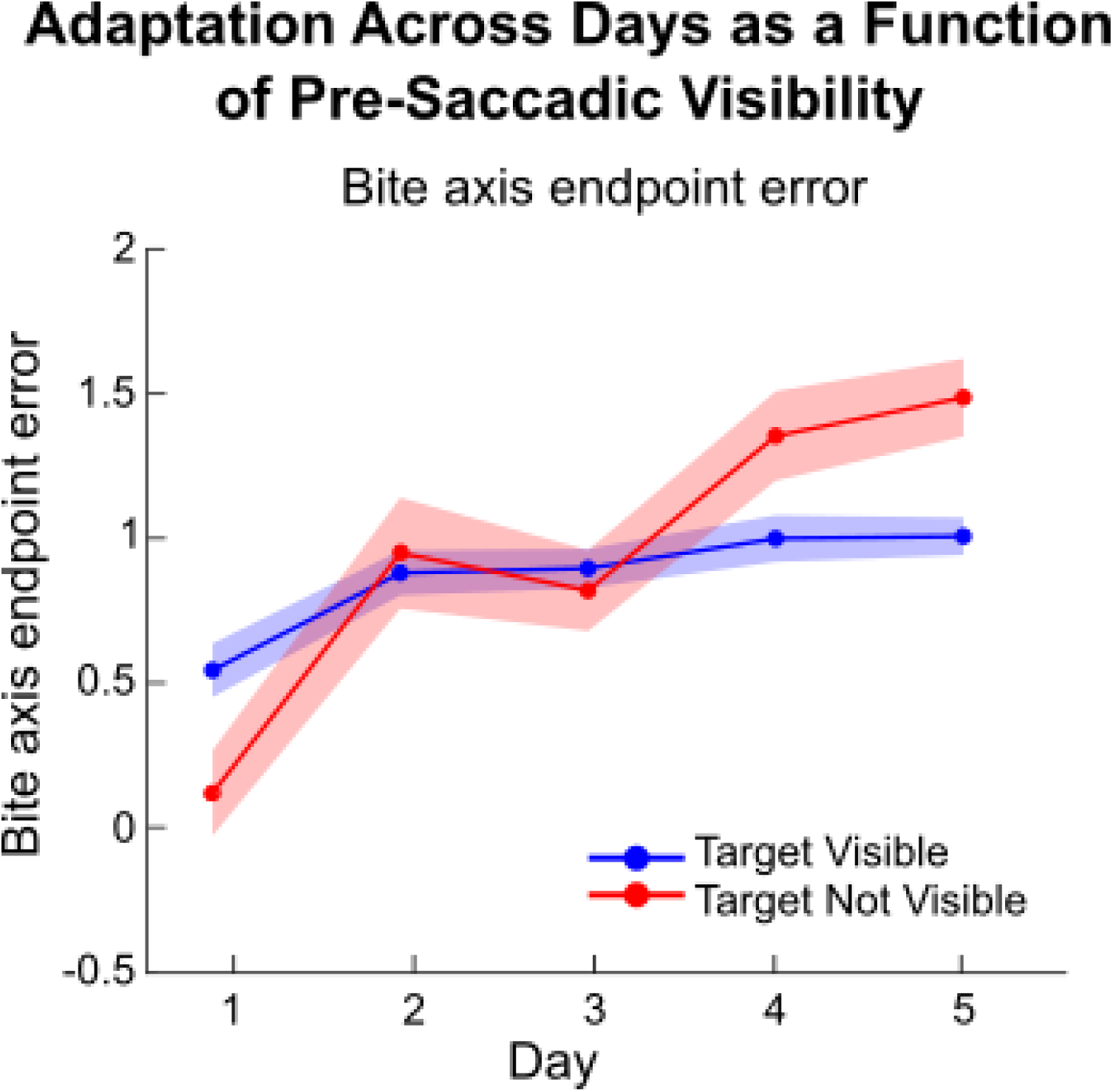
Adaptation across days with and without target visibility. Comparison of bite-axis endpoint errors split by post-step visibility across testing days. Solid lines indicate participant weighted medians (blue: target visible; red: target non-visible). Shaded regions represent within-subject standard error of the mean (SEM).

When the two visibility conditions were collapsed, Holm-Bonferroni corrected significant increases in endpoint error were observed between Day 1 and Days 2–5, as well as between Days 2 and 5 and Days 3 and 4-5 (all p ≤ 0.040). No significant differences were observed between Days 2 and 3, Days 2 and 4, or Days 4 and 5 following corrections.

In general, our results suggest that if a participant can see the target before the saccade, it is more likely that they will make a saccade to the PRL, and actually see the target at saccade landing. However, increases in endpoint error are present in both presaccadic visibility conditions. At the beginning of the experiment, there seems to be an advantage to pre-saccadic target visibility, however by the end, there is still improvement in both metrics regardless of pre-saccadic visibility. As seen in figure 12, the bite axis endpoint errors on day 5 are actually higher when the target is occluded before the saccade compared to visible, however this could be due to much larger saccades with less accuracy. It is possible that participants developed a strategy -when the target is not visible, they know to make a saccade in the movement direction and will most likely see the target again. Therefore they could be overshooting into the PRL direction with less accuracy. In general, target visibility before the saccade had a small effect on post saccadic bite axis endpoint errors, but overall increases were still present in both groups from day 1 to 5.

Finally, we wanted to examine how these post-saccadic endpoint errors change as a function of latency, and whether this relationship differs depending on pre-saccadic target visibility. Since latency is thought to reflect the certainty of the decision to execute a saccade, this analysis allows us to probe not just where saccades land, but how visibility affects the speed-accuracy trade-off underlying catch-up saccade triggering. As seen in figure 13, when the target is visible before the saccade, latencies were shorter for bite axis endpoint errors away from bite (into the scotoma) and longer towards the bite (away from the scotoma, towards the PRL). When the target was not visible before the saccade, latencies were longer, but remained relatively unchanged across endpoint errors. This is consistent with previous studies that find that greater uncertainty, such as with target occlusion, leads to longer saccade latencies (Coutinho et al., 2021; Doré et al., 2024; Nachmani et al., 2020). In order to fully understand how saccade latency affects these errors, we further quantified the effects of target visibility on latency. This will be the focus of the next section.

**Figure 13.**
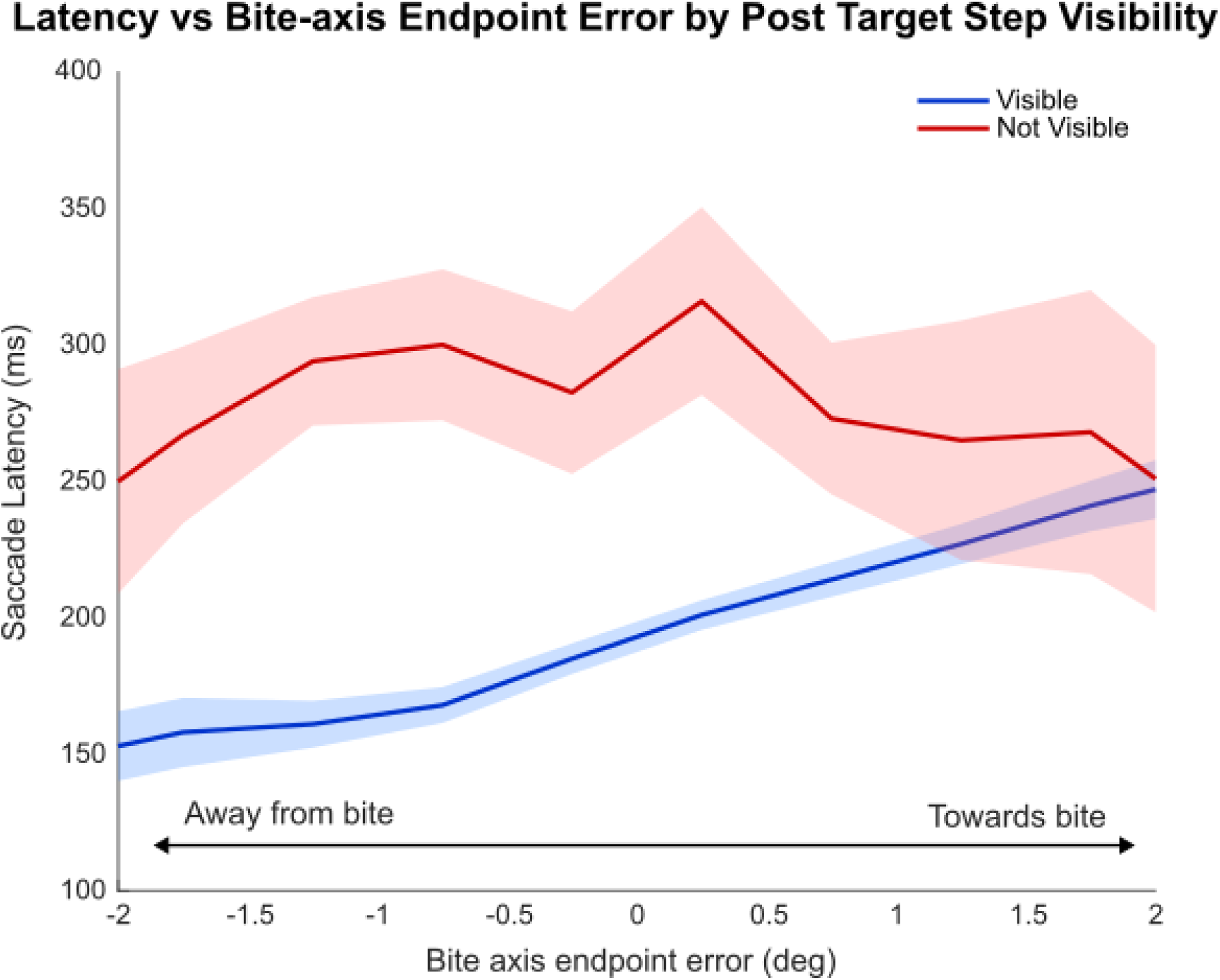
Saccade latency vs. bite axis endpoint error grouped by target visibility. Shaded regions are the 95% confidence intervals around the median saccade latency (solid lines) within each endpoint-error bin of 0.5°.

### 3.6 Saccade Latency and Visibility

To further disentangle the effects of target visibility on saccade timing and potential adaptation, we compared saccade latencies for different visibility conditions on the first and last day of training. We wanted to examine whether saccade latencies changed with different combinations of target visibility at the time of the step, specifically, whether latencies differed with consistent target visibility or occlusion before and after the step, or with target appearance or disappearance after the step. To do this, we compared the participant weighted latencies across 4 groups on the first and last day of training depending on whether the target was visible right before and after the target step (+/-1ms). The participant weighted approach allowed each participant to contribute a single median for each condition and each day. This ensured that each participant contributed equally regardless of the number of valid trials completed in a condition. Figure 14 displays the associated participant medians (panel A) for all conditions and days, as well as violin plots of all participants trials to visualize the differences between days and conditions without participant weighting (panel B). A repeated measures ANOVA comparing the participant weighted median latencies across visibility conditions and days revealed significant main effects of visibility (F(3, 33) = 6.46, *p* = 0.0014559) and day (*F*(1, 11) = 610.41, *p* < 0.0001), as well as a significant interaction effect (*F*(3, 33) = 10.39, *p* < 0.0001). Paired t-tests comparing each condition showed that latency significantly decreased from day 1 to 5 in the visible to visible (p=0.0350) and visible to nonvisible (p= 0.0379) conditions, but changes were not significant for the other two conditions (nonvisible to nonvisible: p=0.5148, nonvisible to visible: p=0.1987). The figures in this section, taken with figure 13, show that between day 1 and 5, saccade latencies generally decrease and lead to increased visibility when the target is visible right before the step.

**Figure 14.**
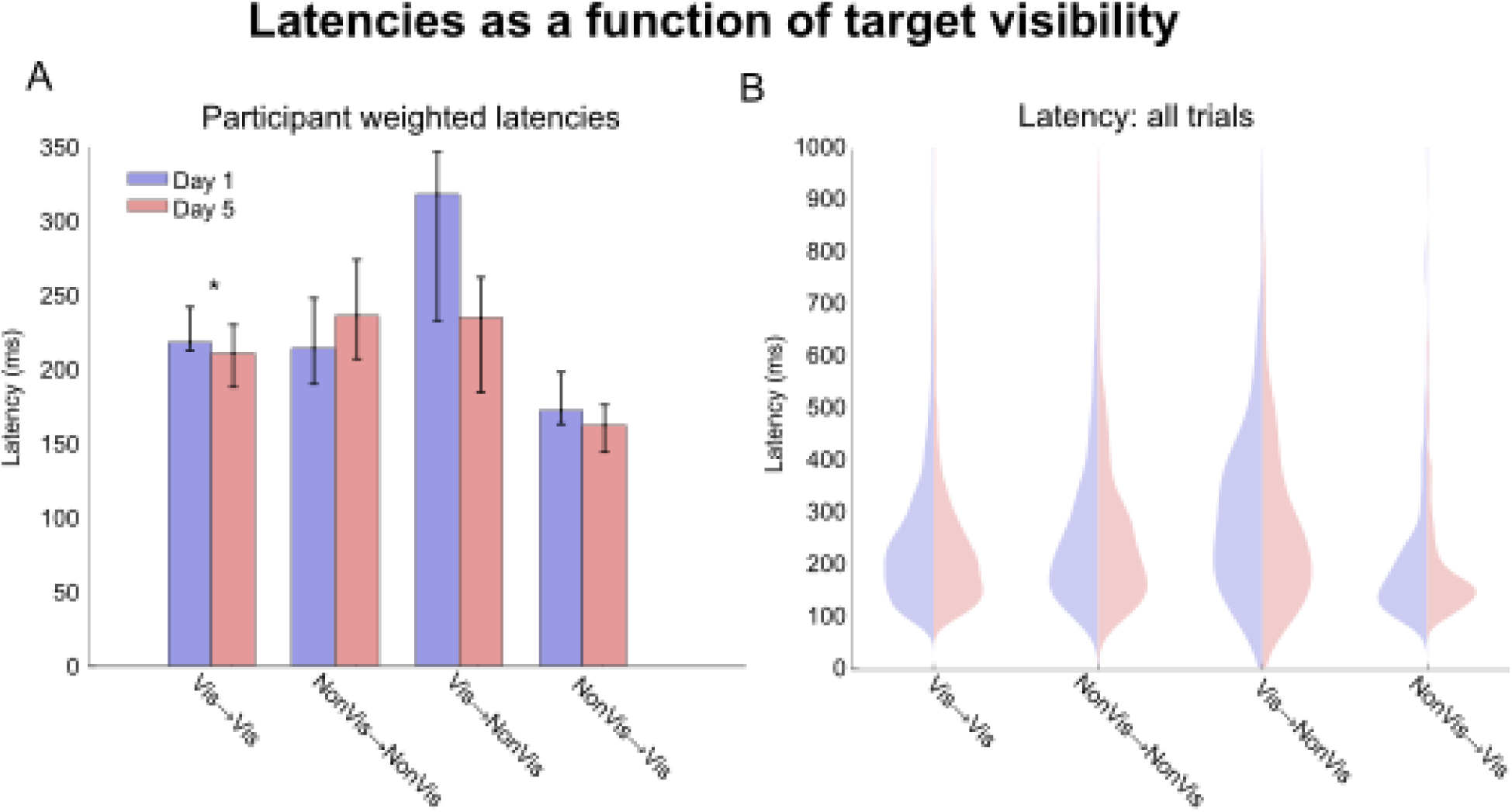
Saccade latencies to target step on day 1 and 5 of training split by target visibility at the step. Panel A displays the participant weighted median latency for each day and visibility condition with the bootstrapped 95% CI of the median as the error bars. Panel B displays the saccade latencies from all trials used in the analysis, not weighted by individual participants. Significance stars from paired t-tests are included for panel A.

### 3.7 Saccade main sequences across control and training days

Saccadic eye movements exhibit a highly conserved kinematic property known as the main sequence, consisting of stereotypical relationships between saccade amplitude, peak velocity and duration, (Bahill A. et al., 1975). This relationship is thought to have evolved as a strategy to optimize the trade-off between speed and accuracy when planning eye movements (Harris & Wolpert, 2006). While saccadic adaptation is not known to change the main sequence relationships in humans (Collins et al., 2008), changes in main sequence relationships have been shown in clinical populations with central field loss, such as those with AMD (Guadron et al., 2023). To examine whether artificial central scotomas, which influence central field visibility, may influence saccades, we examined main sequence relationships for participants’ catch-up saccades without a scotoma during control, and on the first and last day of experimental training. Due to the addition of saccade and pursuit components for catch-up saccades in particular, saccade amplitudes and peak velocities were corrected to remove the pursuit component (de Brouwer et al., 2002) before further analysis (see methods). We then compared linear fits of the main sequence across the control condition, the first day of training and the last day of training. The root mean squared deviation (RMSD) of individual saccades was then computed from each participant and conditions fitted regression. RMSD represents the tightness (or variability) of the amplitude-velocity and amplitude-duration main sequence relationships: lower RMSD reflects more stereotyped, kinematically consistent saccades, while higher RMSD reflects greater trial-to-trial variability relative to the expected main-sequence relationship.

Figure 15 displays the main sequence relationships for a single participant. In this example, we see a slight tightening of the main sequence relationship from day 1 to 5, indicating increased consistency of the kinematic relationship throughout training. For both peak velocity (Panel A) and duration (Panel B) main sequence relationships, this participant shows a decrease in RMSD of the linear fit from day 1 to day 5, reaching a similar level of consistency as the control condition, where there was no scotoma present. For all participants, a repeated measures ANOVA revealed a significant effect of training day (Day 1, 5 or control session) on the RMSDs in the amplitude and peak velocity main sequence, F(2,48)=5.03, p=0.010, indicating that deviations in the main sequence changed across the course of training. In contrast, the RMSDs of the amplitude and duration main sequence did not change over time, F(2,48)=2.90, p=0.065. Paired t-tests for RMSDs for the amplitude and peak velocity main sequence indicated significant differences between control and day 1 (t(24)=-2.84, p=0.027), and day 1 and day 5 (t(24)=2.66, p=0.041) but not control and day 5 (t(24)=-0.81, p=1.000). The increased RMSDs from control to day 1 suggests less consistency in saccade planning once participants initially experience central vision loss. The decrease from day 1 to day 5 and lack of significant differences between day 5 and control suggest more consistent saccade planning over training comparable to baseline behavior with a fovea. These results indicate an ability for participants to adapt the way in which they execute saccades throughout the course of training.

**Figure 15.**
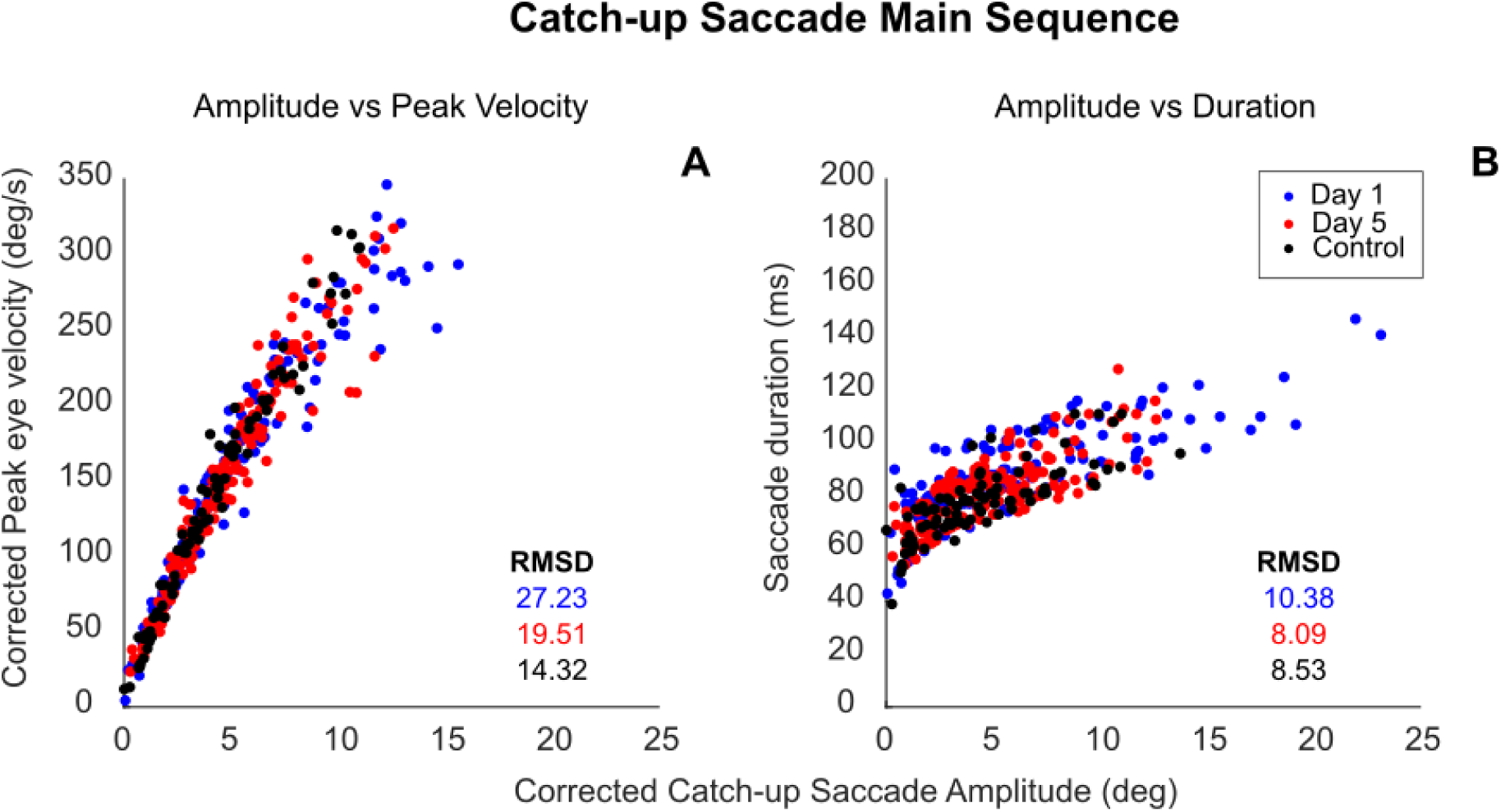
Catch-up saccade main sequence for an individual participant. Catch-up saccade amplitudes and peak velocities have been corrected to remove the pursuit component. Linear fit and root mean squared deviation were computed for each relationship on day 1, day 5 and the control sessions without a scotoma.

## 4. Discussion

### 4.1 General discussion

We investigated whether young, fully sighted participants with gaze contingent scotomas could adopt a preferred retinal locus (PRL) instead of the fovea as their new oculomotor reference for catch-up saccades during smooth pursuit. We observed a shift in catch-up saccade endpoint errors from the fovea to the PRL for most participants, with asymmetries relative to the scotoma’s bite axis, depending on pursuit direction, with larger errors when pursuit motion was into the bite (towards cartesian quadrant 1). In this direction, landing the target onto the PRL requires the oculomotor system to execute larger, overshooting saccades relative to the target’s trajectory, increasing motor variability and residual error. We also observed that saccade latencies varied as a function of target visibility at the time of the target step, with the longest latencies occurring when the target was visible before the step and occluded after, and the shortest latencies when the target was occluded before the step, and visible after. Finally, we found that the root mean squared deviation of the saccadic main sequence decreased over training days, suggesting that training increased the consistency in the catch-up saccade main sequence, with similar variability on day 5 as the baseline in the control task.

### 4.2 Comparison to literature

Our findings are consistent with a growing body of literature demonstrating that adaptation from the fovea to the PRL occurs under both simulated and real scotoma conditions. In simulated scotoma paradigms, Song and colleagues found that saccadic adaptation develops with both visible and invisible gaze-contingent scotomas, concluding that peripheral retinal error is sufficient to drive saccadic adaptation (Song et al., 2021). Similarly, Léné and colleagues reported progressive shifts in saccade endpoints over ten days of practice, indicative of changes in saccade planning and improved eccentric viewing (Léné et al., 2020). Maniglia and colleagues also demonstrated gradual saccadic re-referencing over prolonged training, although, as in the present study, they observed substantial inter-individual variability for learning rates and performance (Maniglia et al., 2020). Kwon and colleagues further showed that shifts of saccade endpoints from the fovea to a PRL during visual search tasks began after only a few hours of training with a simulated scotoma, with complete saccadic referencing occurring within about 15-25 hours of explicit training (Kwon et al., 2013).

Although participants in our study did not receive explicit PRL training, adaptation nevertheless occurred within a shorter time frame of five days. One possible explanation is the design of our simulated scotoma. Unlike the circular or near circular simulated scotomas used in previous studies, our scotoma shape consisted of a directional ‘bite’ that encouraged the development of a PRL in one specific area, rather than having participants adopt arbitrary PRLs elsewhere. We also ensured our scotoma was visible to participants as scotoma awareness has been shown to facilitate saccadic re-referencing for individuals with PRLs (Janssen & Verghese, 2016). Together, these findings suggest that the spatial characteristics and visibility of a simulated scotoma can substantially influence the rate of PRL development.

Evidence from patients with real central vision loss similarly demonstrates oculomotor re-referencing toward a PRL (White & Bedell, 1990) as well as improvements in eccentric viewing with training during fixation tasks (Nilsson et al., 2003). Direct comparisons with these studies, however, should be interpreted cautiously because most previous work has employed visual search or static fixation paradigms, whereas our task required continuous pursuit of moving targets with catch-up saccades. Consequently, our findings extend the existing literature by demonstrating PRL adaptation during dynamic eye movements rather than under static viewing conditions.

Our findings regarding saccade latency are also consistent with recent models proposing that saccades are initiated when accumulated retinal error reaches a confidence threshold. According to these models, greater uncertainty about whether a corrective saccade is required delays saccade initiation, whereas higher certainty produces shorter latencies (Coutinho et al., 2021; Nachmani et al., 2020). In our task, catch-up saccades were triggered by the target step, making target visibility right before and after this step a direct determinant of confidence. When a target transitioned from visible to occluded by the simulated scotoma, this likely reduced confidence that a corrective saccade was necessary, resulting in longer catch-up saccade latencies. Conversely, when the target re-emerged from behind the scotoma, visual error became immediately available, increasing confidence that a corrective saccade was required and producing significantly shorter latencies. These findings suggest that target visibility directly modulates the decision process underlying catch-up saccade initiation. We also examined whether the main sequence of catch-up saccades changed during adaptation to simulated central vision loss. Previous work in patients with age-related macular degeneration has shown that saccade dynamics depend on target visibility at the time the movement is initiated (Guadron et al., 2023). This may explain the increased variability in the main sequence observed at the beginning of training, when participants were still adapting to the simulated scotoma and target visibility was less stable. As participants developed a functional PRL and became more proficient at tracking the moving target, target visibility likely improved, leading to more consistent saccade generation. Consequently, main sequence variability progressively decreased and approached the values observed during control trials, suggesting that oculomotor adaptation restores more stable saccade programming despite the absence of foveal vision.

### 4.3 Hypothetical Neural Mechanisms

Our results show that healthy participants rapidly adapted to simulated central vision loss by shifting their saccade endpoints toward a preferred retinal locus (PRL), demonstrating oculomotor re-referencing, and reducing the variability of catch-up saccades over time. Together, these findings suggest that the oculomotor system can quickly establish a new reference point for guiding eye movements, even in the absence of retinal damage.

The neural mechanisms underlying this adaptation remain speculative. One possibility is that repeated use of an eccentric retinal location gradually changes how visual information is linked to eye movement planning. Instead of using the fovea as the reference for fixation, pursuit, and saccades, the brain may begin using the PRL. This could explain the progressive shift in saccade endpoints and the improvements in pursuit that we observed over the course of the experiment.

The use of a PRL as an oculomotor reference may involve the superior colliculus (SC), given its central role in saccade planning and target selection. Specifically, neurons in the SC respond to information about target position, and position errors. The SC also provides position-related signals to the smooth pursuit system in addition to saccades, which could support the oculomotor referencing of catch-up saccades found in our study (Basso et al., 2000; Krauzlis et al., 1997, 2000; Sparks & Mays, 1990). SC neurons encode information related to target location and position error, providing a link between visual information and the motor commands required to direct the eyes toward a target. Repeated use of a PRL may therefore involve changes in the visuomotor transformations within the SC and its associated circuitry, such that saccade direction and amplitude are specified relative to the PRL rather than the fovea.

These changes likely also involve brain regions responsible for visuomotor control. Previous imaging studies have shown increased activation in frontal and posterior parietal regions during pursuit and saccade tasks in patients with AMD, suggesting greater reliance on attentional and oculomotor networks following central vision loss (Little et al., 2008). In particular, the posterior parietal cortex plays an important role in transforming visual information into motor commands and could therefore contribute to establishing the PRL as a new oculomotor reference. The results of our study suggests that these changes can occur over as little as 5 days, allowing catch-up saccades to be re-referenced to a PRL.

### 4.4 Clinical Implications

Our findings provide a promising direction for rehabilitation strategies aimed at improving PRL use and oculomotor re-referencing in individuals with central vision loss due to macular degeneration. The rapid adaptation observed in our study suggests that targeted eccentric viewing training may help individuals learn to use a PRL more effectively and re-reference eye movements accordingly. Training that combines smooth pursuit with saccadic eye movements may be particularly useful for developing more consistent PRL referenced eye movements. Further studies examining individuals with macular degeneration is needed to determine whether these adaptations can be induced in patients and whether they translate to improvements in visual function.

### 4.5 Limitations

While our experiment provides insight into PRL adaptation during visual tracking, an important limitation is that our participants were young adults with healthy vision rather than patients with AMD. Although this limits the direct clinical relevance of these findings, testing young fully-sighted participants allowed us to determine whether rapid oculomotor re-referencing to a PRL is possible and suggests designs for future paradigms with older adults or patients with AMD.

Another limitation is that simulated scotomas cannot fully replicate the characteristics of real central vision loss. Scotomas associated with AMD may develop only in one eye and vary in size, shape and retinal location, whereas our simulated scotoma was a fixed shape specifically designed to encourage PRL location in a predetermined location. Thus, comparison between real and simulated scotomas is difficult. Whether the same rate and extent of oculomotor re-referencing to a PRL occur in patients with real scotomas is still to be determined.

Finally, participants were not given explicit instructions on how to develop or re-reference to a PRL. Instead, they were simply instructed to track the target to the best of their abilities and were free to explore different strategies. Nevertheless, most participants developed a functional PRL over the course of the experiment with oculomotor re-referencing. Future experiments with older or clinical populations may benefit from more explicit PRL training. For example, one an optimal PRL has been identified, providing explicit feedback using verbal instructions and a gaze contingent display of the PRL and scotoma borders could facilitate faster and more consistent adaptation to oculomotor referencing.

### 4.6 Conclusions

Our study provides evidence that young, fully sighted individuals can rapidly adapt to a gaze contingent central scotoma and use a PRL in place of their fovea as a new oculomotor reference during catch-up saccades. Most participants showed measurable changes in oculomotor referencing after only five hours of exposure to the simulated scotoma, highlighting the plasticity of the oculomotor system. These findings provide further evidence that PRL development can occur over relatively short timescales and extend previous work by demonstrating adaptation during continuous visual tracking rather than static fixation or visual search. Although these results must be confirmed in patients with AMD, they support the potential for rehabilitation strategies that promote efficient PRL use to improve oculomotor function and optimize the use of residual vision in individuals with central vision loss.

## Data Availability Statement

Anonymized data for this project has been made openly available: https://osf.io/fka6u/overview.

## Credit Contribution

**Sydney Doré:** Project administration, Formal analysis, Data curation, Writing – Original draft, Visualization, Investigation. **Jenna Black:** Data curation. **Gunnar Blohm:** Funding acquisition, Methodology, Software, Writing – Review and Editing, Supervision, Resources. **Aarlenne Z Khan:** Conceptualization, Funding acquisition, Methodology, Software, Writing – Review and Editing, Supervision, Resources.

## Acknowledgements

Thank you to the members of the Blohm Lab for valuable project feedback.

## Abbreviations

AMD,: Age-related Macular Degeneration
PRL,: Preferred Retinal Locus
SC,: Superior Colliculus
TRL,: Trained Retinal Locus;

